# Widespread remodeling of the RNA editome underlies transcriptional and clinical heterogeneity in pediatric acute lymphoblastic leukemia

**DOI:** 10.64898/2025.12.03.692208

**Authors:** Sara Brin Rosenthal, Thomas Whisenant, Roman Sasik, Maria Rivera, Geena Il Defonso, Charles-Alexandre Roy, Sabina Enlund, Rawan Shraim, Kathleen Fisch, David T Teachey, Frida Holm, Dennis John Kuo, Qingfei Jiang

**Affiliations:** Center for Computational Biology & Bioinformatics (CCBB), University of California, San Diego, La Jolla, 92093-0681; Division of Regenerative Medicine, Department of Medicine, University of California, San Diego, La Jolla, CA; Moores Cancer Center, University of California, San Diego, La Jolla, CA 92037; Department of Women’s and Children’s Health, Division of Pediatric Oncology and Pediatric Surgery, Karolinska Institutet, Sweden; Perelman School of Medicine, University of Pennsylvania, Philadelphia, PA; Division of Oncology and Center for Childhood Cancer Research, Children’s Hospital of Philadelphia, Philadelphia, PA; and Department of Medicine, Stanford University, Stanford, CA; Department of Neuroscience, Karolinska Institutet, 171 77 Stockholm, Sweden; Division of Pediatric Hematology-Oncology, Rady Children’s Hospital San Diego, University of California, San Diego, CA

## Abstract

Aberrant RNA editing by adenosine deaminases is increasingly recognized as a source of transcriptional diversity in adult cancer, yet its role in pediatric leukemia remains poorly understood. Here, we systematically profiled RNA editing events from bulk RNA-seq data of 1,025 pediatric T-cell acute lymphoblastic leukemia (T-ALL) samples, 37 matched B-ALL diagnosis/relapse pairs, and 6 age-matched non-leukemic controls. We uncovered a widespread global increase in A-to-I editing in T-ALL, affecting both non-coding Alu elements and coding sequences. By contrast, B-ALL shows relatively modest increases in RNA editing at both diagnosis and relapse with no differences in ADAR1 levels. ADAR1 expression is strongly associated with double-stranded RNA (dsRNA) sensors and interferon signaling in T-ALL. Genelevel analyses highlight recurrent editing of oncogenic drivers and regulators of immune signaling, chromatin remodeling, and RNA processing. Unexpectedly, increased editing levels in select genes (*CD247*, *PRKCA* and *TRAF3IP2.ASI* in T-ALL; *SPPL3* in B-ALL) were significantly associated with better patient survival, suggesting a potential prognostic role for editing dysregulation at individual gene levels. Together, these results deepen our understanding of the pediatric ALL transcriptome landscape and provided novel candidate regulators and therapeutic targets for future mechanistic and translational investigation.

## Introduction

Acute lymphoblastic leukemia (ALL) is the most common pediatric cancer, accounting for approximately 25% of childhood malignancies. The majority of ALL cases are of B-cell lineage (B-ALL), which constitutes about 85% of pediatric ALL cases. By contrast, T-cell ALL (T-ALL), a high-risk subtype comprising ∼15% of ALL, is characterized by aggressive proliferation of immature thymocytes and a distinct transcriptional and mutational landscape^1, 2, 3, 4^. While genomic and transcriptomic studies have uncovered numerous driver mutations and disrupted regulatory networks ^5, 6^, the role of the epitranscriptome in leukemia pathogenesis is not well understood.

Epitranscriptomic RNA modifications act as an additional layer of gene regulation on top of genetic and epigenetic regulation exerted at the DNA level. To date, over 200 different types of RNA modifications have been reported in various cellular systems ^7, 8^. Among them, adenosine-to-inosine (A-to-I) RNA editing is one of the most prevalent RNA modifications in mammals. In this process, the adenosine bases in double-stranded RNAs (dsRNA) are deaminated by adenosine deaminases acting on RNA (ADAR1) enzymes, and subsequently interpreted as guanosine (G) during base pairing. These alternations have a wide range of effects on RNA biology including mRNA recoding, splicing, RNA stability and degradation, miRNA biogenesis, and translational control^9, 10, 11, 12, 13^. A-to-I editing is particularly abundant in primate-specific Alu elements located in introns and untranslated regions, where it influences transcript structure and processing^14, 15, 16^. Since inverted oriented Alu sequences form stable dsRNA structures^17, 18^, A-to-I RNA editing of Alu also serves as a molecular marker to distinguish self-from non-self dsRNA, thereby preventing aberrant activation of the innate immune responses^19, 20, 21, 22^.

Malignant ADAR1 activation and transcriptome-wide over-editing have been extensively studies in many adult hematologic malignancies^23, 24, 25, 26^, but has not been systematically investigated in pediatric cancer. Instead of aging-associated accumulation of somatic mutations, pediatric cancers are driven primarily by developmental and epigenetic defects and typically fewer genetic mutations^27^. Relapse-associated A-to-I editing landscape has been reported in pediatric T-ALL ^28^. In this study, we present a comprehensive analysis of A-to-I RNA editing in 1,025 T-ALL and 37 paired B-ALL patient samples at diagnosis and relapse using RNA sequencing data from the Gabriella Miller Kids First ^29^ and National Cancer Institute Therapeutically Applicable Research to Generate Effective Treatments (TARGETS) cohorts. We compare editing profiles to normal pediatric blood samples, identify genes and regions with differential editing, and integrate these findings with gene expression and clinical outcome data. Our results reveal pervasive and distinct RNA editing patterns in T-ALL and B-ALL, implicating editing as a key layer of transcriptomic dysregulation with potential biological and clinical relevance.

## Results

### Elevated and heterogeneous RNA editing landscape in pediatric ALL

We identified A-to-I RNA editing changes in RNA sequencing data from 1,025 pediatric T-ALL patients had enrolled in Children’s Oncology Group trial AALL0434 from the Gabriella Miller Kids First^29^ initiative (Figure 1 and Table S1). Since ADAR1 is associated with T-ALL relapse^28^, we also included 37 matched diagnosis and relapsed pediatric B-ALL from NCI TARGET (37 matched diagnosis-relapse pairs) to explore if RNA editing is important for B-ALL initiation and relapse. The overall survival, relapse status, and demographic data were available for us to assess the relationship between RNA editing and these clinical risk factors (Table S1). We leveraged peripheral blood samples from pediatric Wilms tumor patients (n = 6) as age-matched non-leukemic controls, as no healthy pediatric blood RNA-seq datasets^30^ were available. This choice ensured age-matched hematopoietic background profiles for comparison, avoiding confounding from adult samples where RNA editing and target gene expression are age-dependent^26^. Because the T-ALL cohort is substantially larger than the control and B-ALL cohorts, downstream analyses incorporated strategies to account for this sample size imbalance.

**Figure 1:**
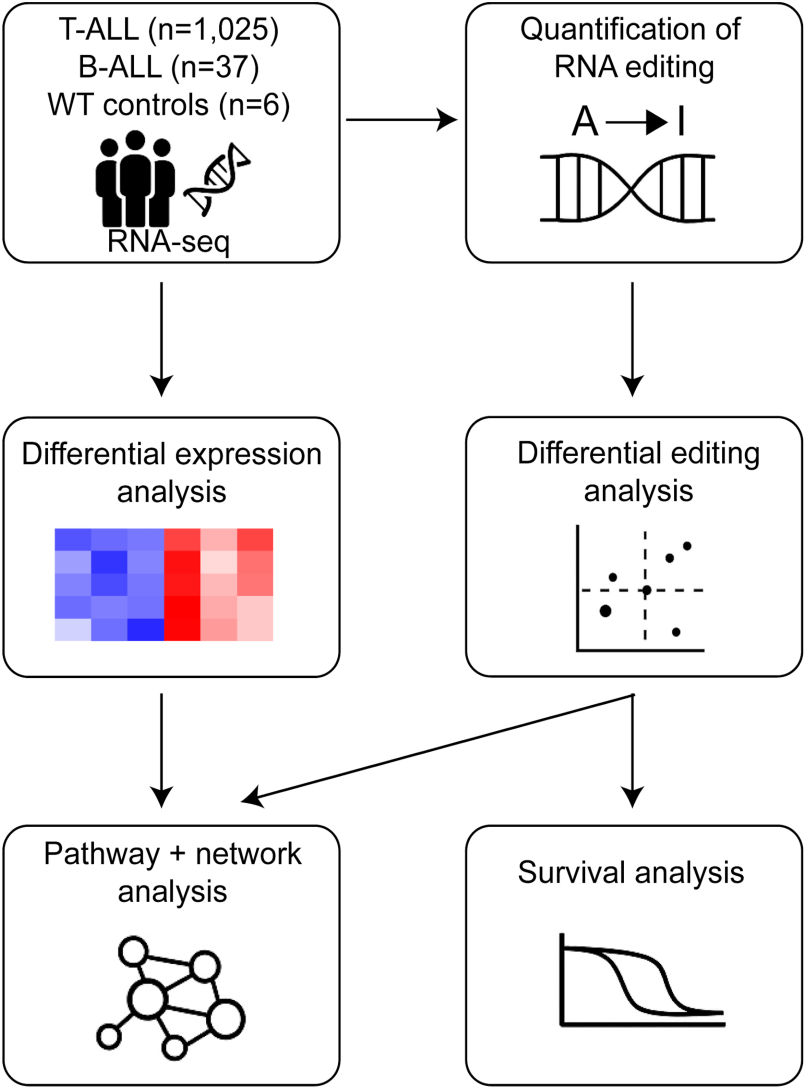
Overview of study design and analysis. This study leverages a large set of T-ALL (n = 1,025), 37 matched B-ALL samples at diagnosis and relapse (n = 37) and age-matched non-leukemia controls from pediatric Wilms tumor patient (n = 6). Comprehensive analyses were conducted to decipher the global RNA editing landscape, individual RNA editing sites, and molecular pathways targeted by ADAR1, and survival analysis.

We first explored global RNA editing patterns across non-leukemic controls, B-ALL (at both diagnosis and relapse), and T-ALL samples (Figure S1). A UMAP dimensionality reduction of editing profiles demonstrated clear separation between non-leukemic controls and T-ALL (Figure 2A). The separation between B-ALL and controls were less distinct, with 17 B-ALL samples clustering near the non-leukemic controls and 20 B-ALL samples clustering near the T-ALL samples. We also do not observe any distinct clusters of B-ALL at diagnosis or relapse. These data point to heterogeneity in editing profiles between T-ALL and B-ALL, with some B-ALL cases resembling normal controls. We also investigated if age, sex or race would contribute to the heterogenous RNA editing landscape and did not find any significant differences among these demographic factors (Figure S2).

**Figure 2:**
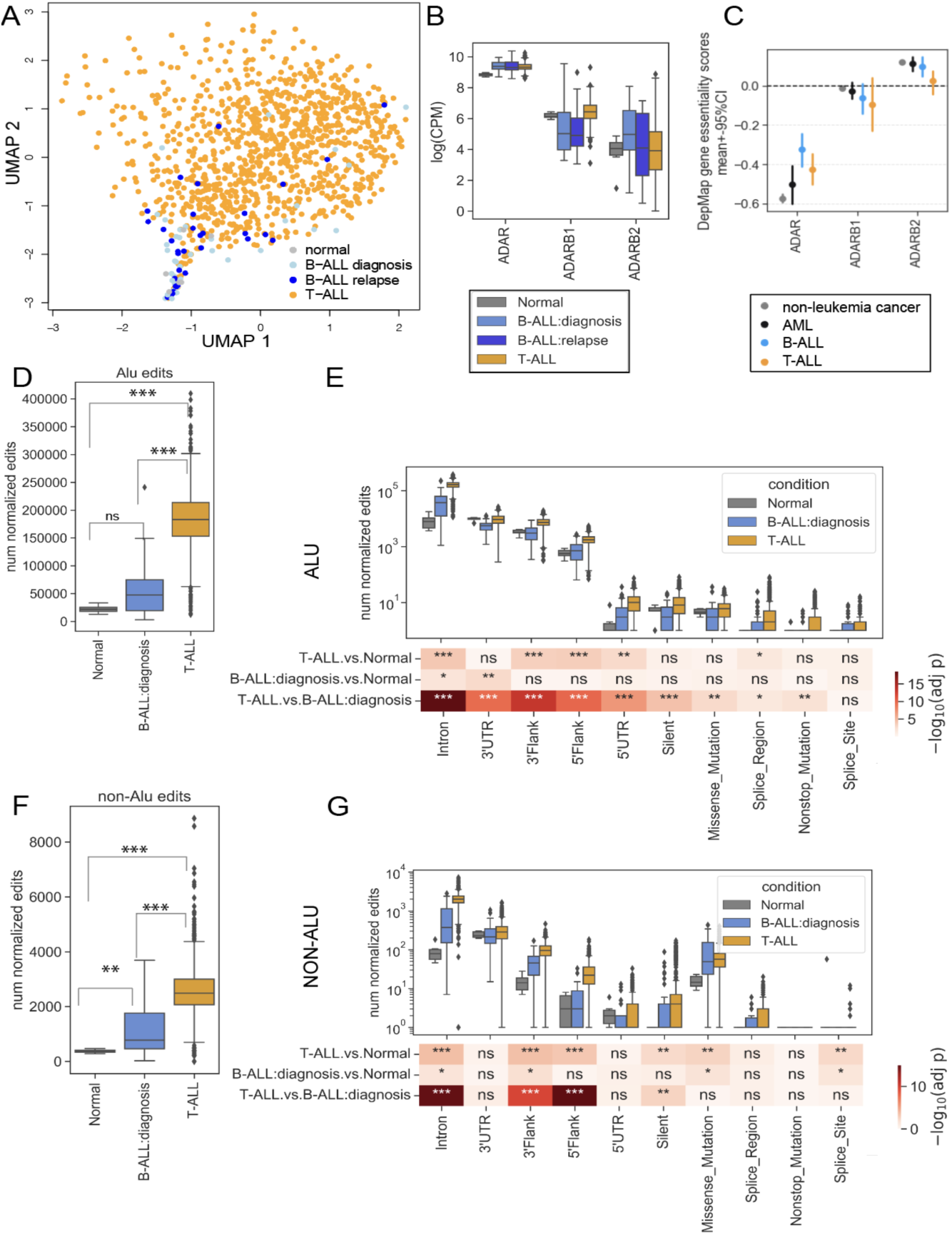
Global RNA editing profiles distinguish T-ALL from B-ALL and normal controls. **A.** UMAP projection of RNA editing profiles derived from normalized editing counts across all samples, colored by condition (T-ALL, n = 1,025; B-ALL, n = 37; **non-leukemic** controls, n = 6). **B.** Boxplot showing expression of ADAR genes across disease types. **C.** DepMap gene essentiality scores for ADAR (ADAR1), ADARB1 (ADAR2), and ADARB2 (ADAR3) across leukemia and non-leukemia cell lines. Lower scores indicate stronger essentiality. Error bars denote 95% confidence intervals. **D.** Boxplot depicting the distribution of total normalized A-to-I editing events occurring in Alu elements across the three conditions. **E.** Distribution of normalized Alu editing counts across annotated genomic features, including intronic, UTR, flank, silent, missense, and splice site categories. A heatmap below the boxplots displays the significance of pairwise comparisons across groups. **F.** Boxplot showing the distribution of normalized A-to-I editing events occurring outside of Alu-regions (non-Alu) across the three conditions. **G.** Distribution of normalized non-Alu editing counts across the same set of genomic features, with corresponding statistical comparisons shown in the heatmap below. ** p<0.01, *** p<0.001, rank sum tests.

We conducted an investigation of the ADAR enzyme responsible for the difference in RNA editing profiles between ALL and normal controls. ADAR1 (*ADAR*) was previously reported as the main RNA editor in hematologic malignancies^25, 26, 28, 31^. In contrast, ADAR2 (*ADARB1*) is mainly expressed in brain tissues and predominately induces protein re-coding editing events^32, 33, 34^. ADAR3 (*ADARB2*) is also specific to brain but so far has no reported A-to-I activity; rather it is considered as a negative regulator of ADAR1 and ADAR2^35, 36^. Indeed, ADAR1 is the highest expressed adenosine deaminase, followed by ADAR2 and ADAR3 (Figure 2B). ADAR1 is significantly overexpressed in T-ALL compared to controls (lfdr = 0.005) and showed a mild increase in expression in B-ALL diagnosis after multiple-test correction (p = 0.0002; lfdr = 0.5). No significant difference was reported in ADAR2 and ADAR3 expression, confirming that ADAR1 is responsible for the differential A-to-I RNA editing landscape in ALL. Furthermore, comparison between matched diagnostic and relapsed B-ALL revealed no significant differences in the expression of any ADAR enzymes. Using the DepMap cell line dataset^37^, we analyzed the gene essentiality score of ADARs for survival of non-leukemia cancer cells, acute myeloid leukemia (AML), B-ALL and T-ALL (Figure 2C). In DepMap genome wide CRISPR screening, essentiality scores quantify the impact of gene loss on cell viability, thus lower scores indicate stronger dependence^37^. ADAR1 exhibits low essentiality scores across different cancer types, suggesting that ADAR1 plays an essential role in both leukemia and non-leukemia cancer. In contrast, the scores for ADAR2 were close to 0 and ADAR3 showed a positive score, which indicates that both ADAR2 and ADAR3 are largely non-essential for leukemia or other cancer types.

We next quantified total A-to-I editing events after normalizing for library size in our sample cohorts (Figure S3). Consistent with previous studies^38, 39^, editing occurred predominantly within Alu regions (98% in normal, 99% in T-ALL, 97% in B-ALL), compared to non-Alu regions (2% in normal, 1% in T-ALL, 3% in B-ALL) (Figures 2D-G). The RNA editing within Alu elements was significantly elevated in T-ALL compared to both B-ALL (p = 6E-24) and non-leukemic controls (p = 1.5E-8), averaging 184,563 high-confident edit sites per sample in T-ALL versus 46,463 in BALL and 22,632 in controls (Figures 2D-E). A similar trend was observed for non-Alu edits, with T-ALL showing the highest levels (2,577 edits per sample), followed by B-ALL (1,162 edits per sample), and controls (374 edits per sample) (Figures 2F-G). In non-Alu edits, T-ALL had significantly increased editing levels compared to both B-ALL (p = 5E-14) and controls (p = 2.5E8). While B-ALL and controls did not have significantly different levels of editing in Alu regions, RNA editing was significantly elevated in non-Alu regions in B-ALL compared to controls (p = 0.01).

Building on these findings, we carefully examined the distribution of editing events across specific variant types within RNA structure. Since the overwhelming majority of RNA editing occurs in Alu-rich regions, we compared the editing pattern in Alu and non-Alu separately (Figures 2E and 2G). T-ALL samples consistently exhibited strong upregulation of Alu editing in introns and regulatory regions (3’ and 5’ flanks, 3’-UTRs), as well as increased editing in silent, missense, and splice-related sites compared to both B-ALL and controls (Figure 2E). By contrast, B-ALL had enriched RNA editing in Alu-containing introns and splice regions compared to controls, while the editing pattern was slightly reduced in 3’UTR and missense positions. A similar locational preference was observed in non-Alu regions between T-ALL and control, although present at lower overall levels (Figure 2G). When comparing B-ALL non-Alu editing to that of controls, we observed significantly elevated editing in introns, 3’ flank, and missense mutations. Interestingly, missense editing was relatively higher in non-Alu regions than Alu-regions in normal controls, B-ALL and T-ALL, which likely reflects the inherent preference of Alu elements within non-coding regions^40, 41^. These results suggest that T-ALL is associated with widespread upregulation of Alu and non-Alu RNA editing, particularly in intronic and regulatory regions. By contrast, B-ALL showed less editing than T-ALL and less preference of RNA editing in Alu-enriched regions.

In addition to the number of edited sites, we also examined the variant allele frequency (VAF), defined as the percentage of edited base at each site (Figure S4). The VAF was not significantly different across conditions, at the level of overall VAF per sample (Figures S4A-B) or the level of individual variant type (Figures S4C-D). Overall, these results point toward shifts in the number of editing sites and position of specific genes and sites, rather than changes of editing level at individual sites.

### Interferon signaling is associated with increased ADAR1 expression in T-ALL, but not overall RNA editing

Interestingly, we observe a striking heterogeneity in the ADAR1 expression and RNA editing levels within the T-ALL cohort. This suggests that RNA editing alterations are not uniformly present across all leukemia cases and may reflect underlying genetic subtypes, variation in ADAR1 expression and activity, or patient-specific regulatory mechanisms influencing editing in leukemic cells. Leveraging the large sample size of the T-ALL cohort, we first examined the potential regulation of ADAR1 expression and pathways influenced by ADAR1 by correlating individual gene expression with ADAR1 expression in T-ALL (Figure 3 and Table S2). ADAR1induced RNA editing collectively mark endogenous dsRNA with inosine bases, which prevents detection by dsRNA sensors such as MDA5, PKR, RIG-I and more recently Za RNA sensor ZBP1 (Figure 3A) ^13, 19^. We identified a total of 385 genes positively associated with ADAR1 expression (R > 0.25 and adj. *p* < 0.05). These genes were broadly enriched for pathways related to interferon activation, anti-viral response, and innate immune response (Figure 3B). The overall IFN signaling strength, as measured by IFN-stimulated gene (ISG) score ^42^, also strongly correlated with ADAR1 expression (Figure 3C). Indeed, all top 5 positively correlated genes are part of the classic inflammation stress module, including *PARP9*, *APOL6*, *MX1*, and *IFI44L* (Figure 3D). Notably, the most significantly correlated gene with ADAR1 is PKR (*EIF2AK2*), a kinase that inhibits global translation synthesis through eIF2 phosphorylation upon sensing dsRNA^13^. By contrast, MDA5 (*IFIH1*) and RIG-I showed no strong association and *ZBP1* is not expressed in our dataset (Figure 3E), suggesting that a selective IFN-ADAR1-PKR regulatory axis may play dominant role in TALL. Interestingly, although we cannot perform correlative analysis in B-ALL due to the small sample size, several ISGs (*EIF2AK2, MX1,* and *IFI44L*) are strongly correlated with ADAR1 expression in B-ALL regardless of relapse status, implying IFN signaling may be a common driver of ADAR1 expression in pediatric leukemia (Figure S5).

**Figure 3:**
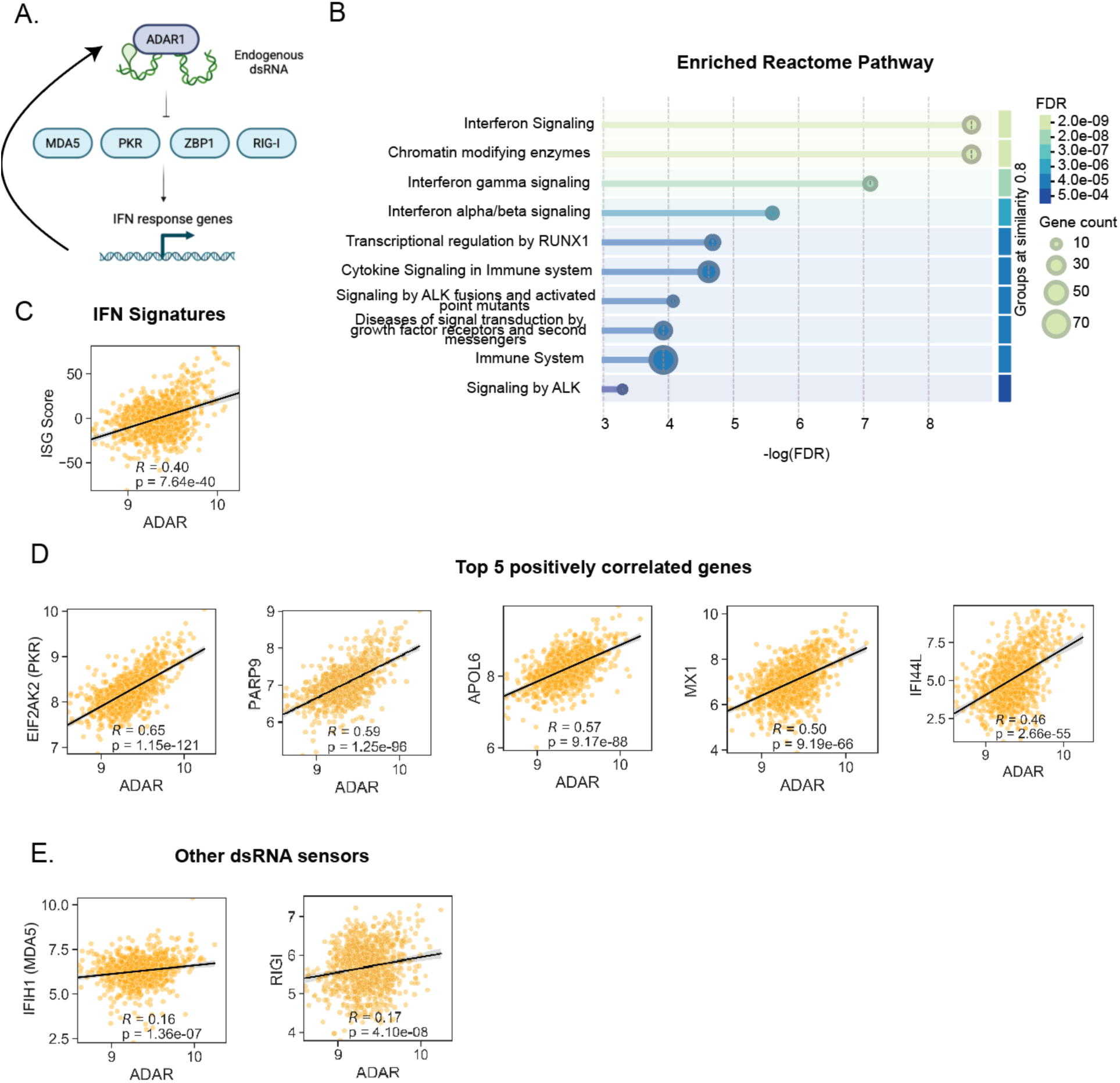
Interferon signaling dictates ADAR1 level in T-ALL. **A.** ADAR1-mediated dsRNA sensing pathways control cellular IFN level. **B.** Pathways analysis of the most significantly correlated genes with ADAR1 expression (R > 0.25 and adj. *p* < 0.25). Multiple pathways are related to inflammation and IFN signaling. **C-E**. Correlation of ISG score^42^ (C), top 5 positively correlated genes (D), and known dsRNA sensor (E) with ADAR1 level.

We next explored the potential regulators of RNA editing by applying a linear regression model to identify genes whose expression correlated with global editing levels in both Alu and non-Alu regions (Figure S6 and Table S3). However, no genes exhibited significantly association with the global RNA editing level, with *ZNF519* showing the highest yet moderate correlation level (R² = 0.089 and p = 1.39E-22; R² = 0.132 and p = 2.79E-33 in Alu and non-Alu, respectively). Pathway analysis of the top 500 genes positively associated with editing revealed enrichment for doublestranded DNA binding and nucleic acid metabolic progress, whereas negatively associations were primarily linked to RNA splicing pathways (Table S3). Surprisingly, ADAR1 expression did not strongly correlated with overall editing level in Alu (R^2^ = 0.004, p = 0.034) or non-Alu sites (R^2^ = 0.018, p = 1.46E-05). Similarly, no significant correlations were observed between RNA editing and ADAR2 or ADAR3. Together, these data indicate ADARs level alone cannot account for the editing patterns in the cellular context of T-ALL.

### Detection of disease-type specific RNA recoding events

Although relatively rare compared to RNA editing in non-coding regions, A-to-I RNA modifications can introduce nonsynonymous changes in protein coding regions^43, 44^. We investigated the most common recoding events in both Alu and non-Alu sites across ALL and normal controls (Figures 4A-B and Figure S7). An average of 10 recoding events per sample in T-ALL, 8 per sample in diagnosis B-ALL, 6 per sample in B-ALL relapse, and 5 per sample in normal controls were reported (Table S4). These events spread across missense, nonstop, or splicing sites. Several genes (e.g. *ZNF506*, *FLNB*, *NEIL1*, and *COG3*) displayed a high level of editing across all sample cohorts, which likely reflects ubiquitous edited sites that are essential in RNA processing rather than leukemia-specific events (Figures 4A-B).

**Figure 4:**
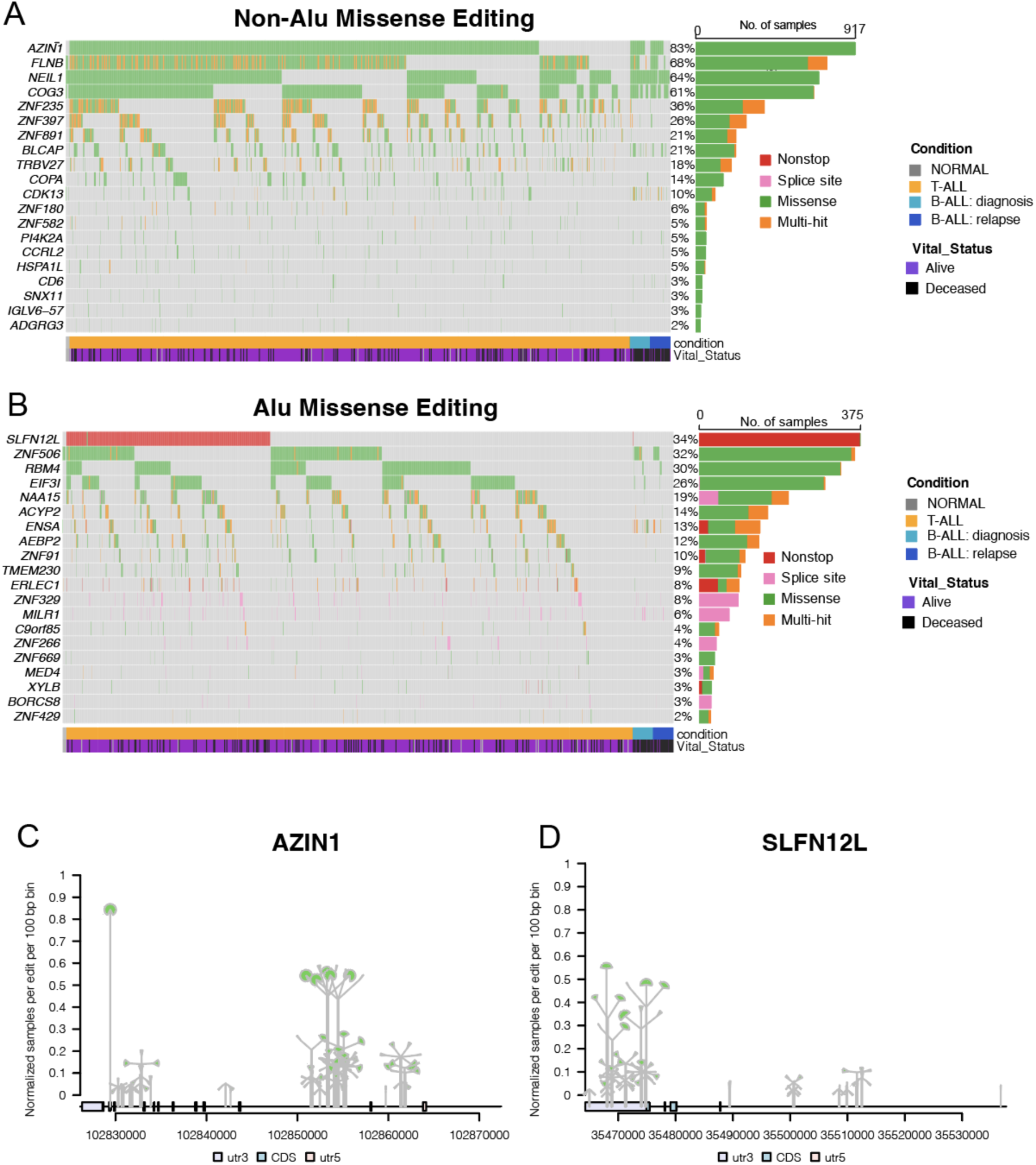
Gene-level RNA editing patterns in coding regions across T-ALL, B-ALL, controls. **A.** Oncoplot of top 20 genes with the largest number of non-Alu coding edits across TALL, B-ALL, and controls. Columns represent samples, and rows represent genes. Color bars to the right indicate the proportion of samples with missense (green), splice site (orange), nonstop (light blue), or multi-hit (black) edits in each gene. **B.** Oncoplot of top 20 Alu-associated coding edits across T-ALL, B-ALL, and controls. Layout and annotations are as in panel (A). **C-D.** Dandelion plots showing editing location within *AZIN1* and *SLFN12L* transcripts in T-ALL. At the top of each stem, a pie chart shows the percentage of individuals with an edit at the site. The yaxis denotes the normalized edit prevalence (fraction of samples edited) for each genomic bin. Gene models are shown below each plot.

Other genes, including *AZIN1*, *SLFN12L*, *RBM4*, and *ZNF235* were significantly more edited in T-ALL and B-ALL compared to controls (lfdr < 0.05). Notably, we identify a canonical S367G missense edit *AZIN1*^45, 46, 47^ with elevated editing levels in T-ALL. We identified missense edits in *AZIN1* in 864 T-ALL (84% of samples) and 28 B-ALL diagnosis (76% of samples), compared to 0 missense edits in normal, which represents a significant increase in both T-ALL (adj. *p* = 7E-6) and B-ALL (adj. *p* = 0.007). Thus, *AZIN1* missense editing likely represents a pan-cancer oncogenic event in both adult and pediatric cancers. We also observed recoding events found only in controls. For example, a missense edit in *DACT3*, a known tumor suppressor with known relevance to leukemia^48^, was detected in 3 normal samples but was not detected in any T-ALL or B-ALL samples (adj. *p* = 3E-111 and 3E-4, respectively) (Figure S7). This striking depletion suggests editing of *DACT3* may act as a protective mechanism that is selectively lost in leukemic transformation.

In order to evaluate the robustness of our results given the small size of the control cohort compared to T-ALL, we performed a sub-sampling analysis. In the subsampling procedure, coding edits were quantified in 25 randomly selected T-ALL samples and compared to the 6 normal controls. Across 100 sub-sampling iterations, the *AZIN1* and *DACT3* missense edits remained significantly different between T-ALL and controls in 99 and 100 runs, respectively (adj. p < 0.05), indicating that the observed differences were not confounded by the sample size imbalance.

In Alu-enriched regions, the top differentially recoding events occurred in *SLFN12L*, *ZNF506* and *RBM4*, all of which were not previously reported in any cancer types (Figure 4B). *SLFN12L* is a member of the Schlafen family which regulates T-cell development and differentiation, as well as immune regulation^49, 50^. Interestingly, the edits are largely located within the 3’ UTR region of *SLFN12L*, which result in non-stop mutations (Figure 4D). We detected a total of 369 nonstop RNA editing events in *SLFN12L* in T-ALL (36% of samples), compared to 2 nonstop edits in BALL diagnosis (5% of samples), and not detected in normal controls. Comparison between *SLFN12L* edits in T-ALL and that in B-ALL suggested these events are T-ALL specific (p.adjust = 4E-3). Similarly, in *RBM4* transcripts, 321 missense edits were reported in T-ALL (31% of samples), compared to 4 in B-ALL (11% of samples), and 0 in normal controls. Other RNA editing targets, such as *ZNF506*, displaced comparable levels of editing among T-ALL, B-ALL and controls (26%, 35%, and 83% of samples, respectively). These findings point to a cell-typespecific missense editing program driven by ADAR1 in T-ALL, characterized by distinct recoding events in key regulatory genes.

### Non-coding differentially edited transcripts between pediatric ALL and normal controls

Since RNA editing predominantly occurs in non-coding regions within the gene body, we next investigated editing at the gene level by integrating signals across all annotated sites within each gene. We used a resampling approach^51^ that repeatedly draws matched sets of T-ALL and control samples after removing the overall disease effect, allowing us to generate fair comparisons despite the unequal group sizes and to obtain well-calibrated p-values and FDR estimates (see Materials and Methods, Figure S8). We find highly significant and widespread differentially edited genes between both T-ALL and B-ALL to controls (Figure S9). A total of 1,605 transcripts exhibits significantly elevated level of editing in T-ALL compared control (FDR < 0.05, coef. T > 10), while only 4 genes showed a lower editing level in T-ALL (FDR < 0.05, coef. T < -10) (Figures S9A-B, Table S5). The strong bias toward increased editing level in T-ALL is consistent with the global pattern of editing (Figure 2) and the previously reported pattern of increased editing levels in leukemia^52, 53, 54^. Since the B-ALL sample size is significantly smaller than T-ALL, a lower stringency was applied to detect differentially edited transcripts between B-ALL and normal controls. We observed 530 genes with higher editing levels (FDR < 0.05, coef. B > 5) and 60 genes with lower editing levels in B-ALL diagnosis compared to control (FDR < 0.05, coef. B < 5) (Figures S9C-D, Table S5).

### Integration of differential editing and gene expression to identify affected biological pathways in T-ALL

To further explore the molecular targets and the biological processes that may be impacted by RNA editing in T-ALL, we combined the editing profiles with the differential gene expression pattern between T-ALL and normal controls. Since sequencing read abundance can lead to overestimation of editing events in highly expressed transcripts, we first correlated the differential gene expression and RNA editing per gene between T-ALL and normal controls (Figure S10). Indeed, a significant correlation was reported between gene expression and level of editing (p < 1E-200, Pearson correlation = 0.645). To account for the read-depth bias in RNA editing analysis, we build a linear model using limma (see Material and Methods) to account for number of edits per gene in the comparisons between T-ALL and normal controls. This method allows adjustments for the level of expression in each gene. Importantly, our method for detecting differential RNA editing levels explicitly controls for underlying RNA expression levels, ensuring that observed editing differences are not simply a byproduct of increased transcript abundance.

With this newly developed pipeline we examined the RNA editing targets with both elevated editing and differentially expressed in T-ALL compared to control. We identified 297 genes with significant upregulation in both expression and editing, and 365 genes with reduced expression coupled with elevated editing levels in T-ALL (FDR < 0.05, coef. T > 10; expression log fold change > 1, expression log fold change < -1, respectively) (Figure 5A). Several increased expressed and edited genes were involved in normal T-cell function, such as activation, TCR receptor signaling, and proliferation (e.g *CD3D, CD247, LCK*, and *PRKCB*)^55, 56, 57, 58^. Other genes in this category include major epigenetic regulators and tumor suppressors and oncogenes frequently mutated in leukemia and other cancer types (e.g *KMT2C, EP300*, *CREBBP*, *SETD2*)^59, 60, 61, 62, 63^, well-known drivers of leukemic transformation (e.g. *MYB, RUNX1*)^64, 65^, splicing and RNA processing factors (e.g. *CELF1, RBM4*)^66, 67^, and TP53 pathways (e.g. *MDM2, MDM4*)^68, 69, 70^. Genes with elevated editing, but lower levels of expression, include mitochondrial function (e.g. *NDUFC1, NDUFS4*)^71^, protein trafficking and transport genes (e.g. *RAB1A, RAB2A, RAB5B, SEC11A*)^72, 73, 74^.

**Figure 5:**
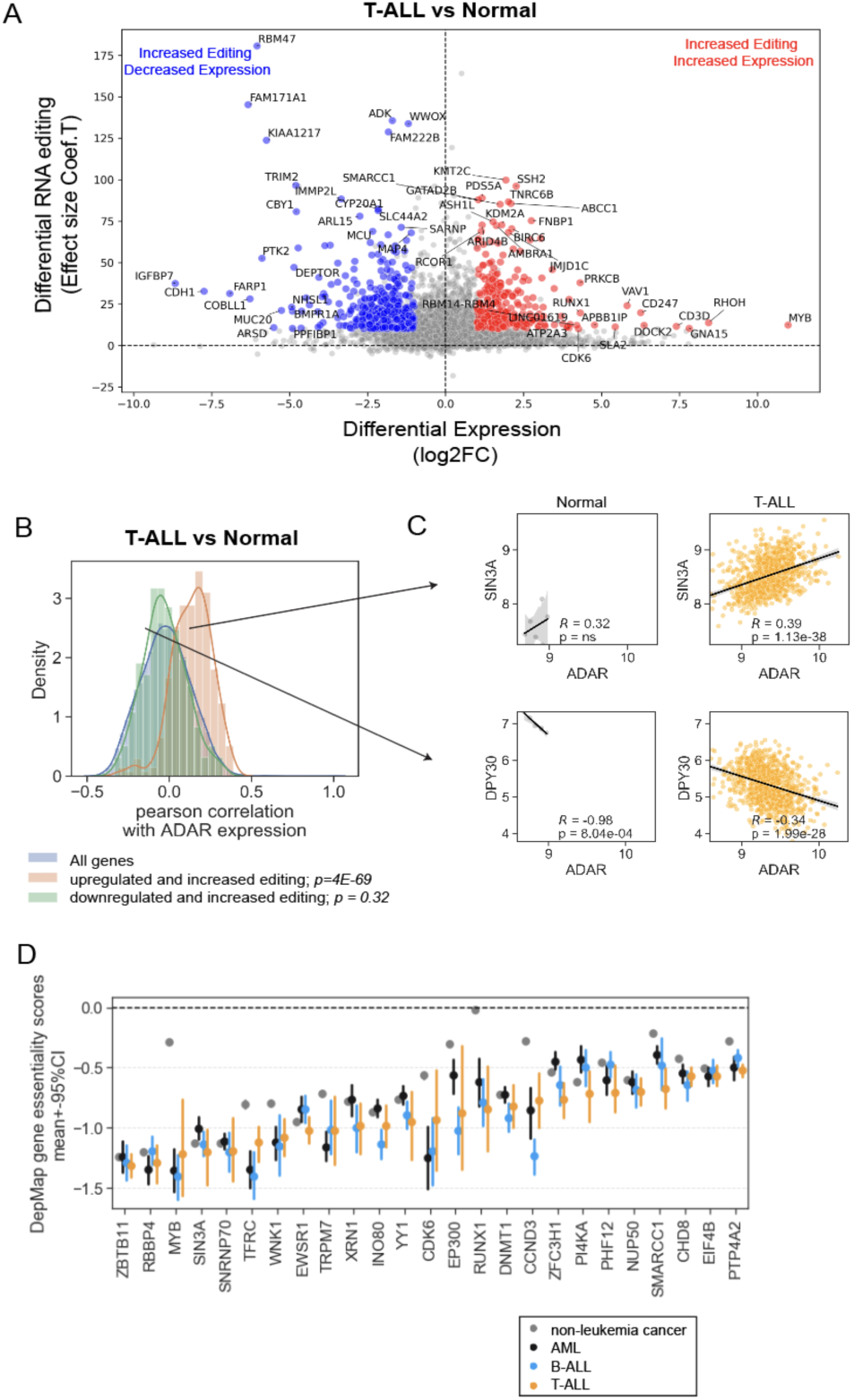
Integration of differentially expressed genes and differentially edited genes in TALL. **A.** Scatterplot showing a positive correlation between log fold change of differentially expressed genes (x-axis) and differentially edited genes (y-axis), between T-ALL and normal control. Red dots indicate concurrently significantly upregulated genes adj. *p* < 0.05 & log FC > 1) and significantly more edited genes (adj. *p* < 0.05, coef. T > 10). Blue dots indicate significantly downregulated genes (adj. *p* < 0.05 & log FC < -1) and significantly more edited Alu-intron regions (adj. *p* < 0.05, coef. T > 10). Select genes are labeled. **B.** Kernel density distribution of Pearson correlation coefficients between ADAR expression and all expressed genes. Genes with both elevated editing and expression in T-ALL (orange) show significantly stronger positive correlation with ADAR compared to baseline (all genes, blue; p = 4E-69, rank-sum test). Genes with elevated editing but reduced expression in T-ALL (green) show lower correlation with ADAR compared to baseline (p = 0.32, rank-sum test), though the effect size is smaller. **C.** Scatterplots showing *SIN3A* and *DPY30* expression significantly positively (most significant of genes in orange peak in panel B) and negatively (most significant of genes green in panel B) correlates with ADAR1. Each point represents an individual sample. The normal control and T-ALL are shown separately. **D.** DepMap gene essentiality scores for selected editing-associated genes across leukemia and nonleukemia cell lines. Lower scores indicate stronger essentiality. Error bars denote 95% confidence intervals.

We then examined the co-dependence of ADAR1 and its potential targets by correlating the overall expression patterns in T-ALL. The upregulated and more edited genes in T-ALL compared to normal displayed a significantly higher co-expression with ADAR1 (Figure 5B, p = 4E-69), while genes that were more highly edited but downregulated exhibited no significant shift, suggesting low expected correlation with ADAR1 (p = 0.32). These findings support the notion that editing-associated upregulated genes are more directly linked to ADAR1 overexpression, where editing-associated downregulated genes may be influenced by ADAR1 through indirect mechanisms rather than an inhibitory mechanism by RNA editing. To confirm this, we correlated individual gene expression with ADAR1 expression in normal control and T-ALL separately. Out of genes with increased editing, the most significantly positively and negatively correlated were *SIN3A* (R = 0.39, P = 1.13E-38) and *DPY30* (R = -0.34, P = 1.99E-28), respectively (Figure 5C).

To explore if any highly edited transcripts have functional relevance, we screened the 294 genes that were both significantly upregulated and more highly edited against DepMap CRISPR essentiality scores. We found 56 out of 297 genes (19%) are essential in T-lymphoblastic leukemia/lymphoma cell lines (score < -0.5; Figure 5D, Table S6). These genes were either pan-essential with strongly negative essentiality scores across all cancer types (e.g., *ZBTB11*, *RBBP4*, *SIN3A*, and *SNRNP70*) or appeared to be leukemia specific with previously established functions in hematological malignancies (e.g. *MYB, TFRC, EP300, CDK6, CCDN3 and RUNX1*) (Figure S11), suggesting RNA editing converges on critical oncogenes in T-ALL.

Further network analysis using the STRING molecular interaction database revealed that both upregulated and downregulated gene sets exhibited significantly more interactions than expected by chance (p < 1E-16) (Figures S12A-B), suggesting that ADAR1 may influence groups of genes involved in shared biological pathways in T-ALL. Gene set enrichment analyses were conducted against known pathway databases including Gene Ontology, REACTOME, KEGG, wikipathways, Pubmed (Figure S12C-D). Although roughly equal numbers of edited genes were up- and downregulated, the pathway enrichments were largely confined to upregulated gene sets. A total of 1,308 biological pathways were identified in genes with elevated expression and editing and 533 pathways were reported in genes with reduced expression and elevated editing (Table S7). Elevated gene sets were broadly enriched for chromatin organization (GO: chromatin organization, FDR = 3.74E-23, Reactome: chromatin modifying enzymes, FDR = 1.10E-17) and leukemia signaling pathways (KEGG: T cell receptor signaling pathway, FDR = 0.0016; GO: cell cycle G1/S phase transition, FDR = 3.68E-07; and Reactome: transcriptional regulation by RUNX1, FDR = 7.08E-09) (Figure S12C). The downregulated gene sets were enriched for intracellular transport, autophagy, endocytosis and membrane trafficking (Figure S12D). Together, these data underscore the potential role of ADAR1 in regulating multiple pathways and contributing to a broad epigenetic and transcriptomic reprogramming in T-ALL.

### Validation of targets in independent ADAR1 knockdown T-ALL datasets

To functionally investigate whether ADAR1 directly regulates important genes and to establish a causal link between RNA editing and differentially expression, we leveraged an independent RNA-seq dataset from T-ALL patient-derived cells modified by either a scramble control or a shRNA targeting ADAR1 to knockdown ADAR1 ^27^. Differentially expressed genes upon ADAR1 knockdown (shADAR1 vs control) were then overlaid with the sets of both differentially expressed and edited genes between T-ALL and non-leukemic control (T-ALL vs normal) (Figures 6A and Table S8). Strikingly, genes with elevated editing and expression in T-ALL were significantly enriched with transcripts downregulated upon ADAR1 knockdown (86 overlapping genes; p = 2E4, Fisher’s exact test). Conversely, genes with elevated editing but reduced expression in T-ALL significantly overlapped with those upregulated by ADAR1 knockdown (83 genes; p = 5E-3, Fisher’s exact test). By contrast, no significant overlapping was observed in the same directional change comparisons. These reciprocal patterns indicate that ADAR1-mediated editing is responsible for driving both activation and repression of target genes in T-ALL.

**Figure 6:**
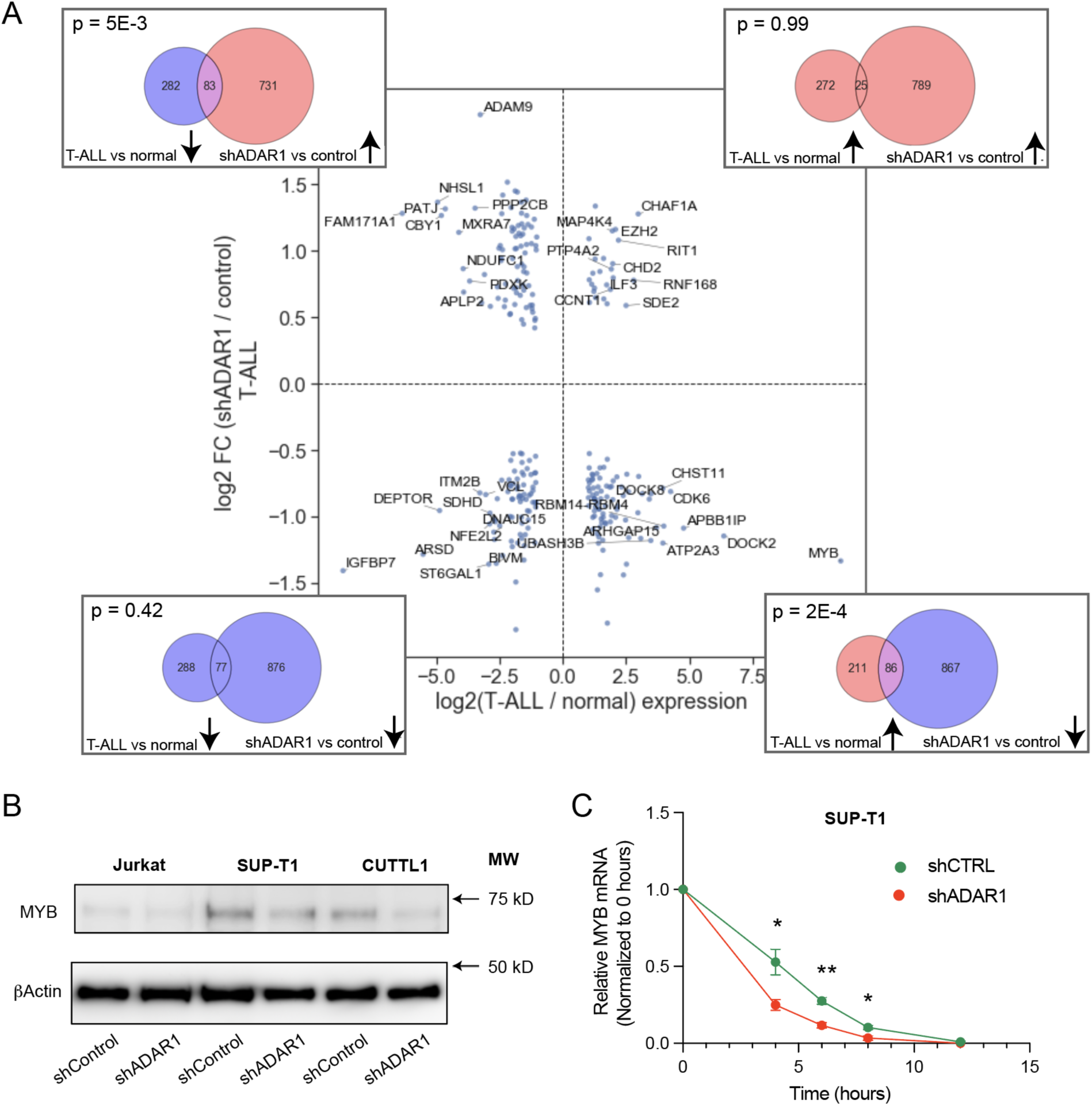
Identification of ADAR1 targets in T-ALL. **A.** Scatterplot showing the relationship between log fold change in expression between T-ALL and normal (x-axis), and log fold change in expression upon ADAR knockdown in T-ALL patient (n = 2) (y-axis). Venn diagrams show the overlap between gene sets in each quadrant, with p values from Fisher’s exact test. **B.** Western blot showing MYB protein level after ADAR1 knockdown in three T-ALL cell lines. **C.** Quantification of remaining *MYB* mRNA level after Actinomycin D treatment in SUP-T1 cells by RT-qPCR. Experimental triplicates, unpaired t-test.

We selectively validated the effect of ADAR1 on MYB, a well-established master transcription factor essential for leukemia proliferation^5, 65, 75^. The T-ALL editing sites in *MYB* clusters to several intronic regions (Figure S11). To investigate the functional relevance of RNA editing on MYB, we performed knockdown studies in three T-ALL cell lines (SUP-T1, Jurkat, CUTTL1) using shRNA targeting ADAR1 (Figure 6B). Indeed, we observed reduction of MYB protein level following ADAR1 knockdown. We further assess the molecular mechanism by measuring the *MYB* mRNA decay following Actinomycin D treatment in SUP-T1 cells which express the highest level of MYB. ADAR1 knockdown significantly accelerates MYB mRNA decay (Figure 6C), suggesting RNA editing in *MYB* intronic region may prevent mRNA degradation. Thus, our results indicate that ADAR1 supports leukemia growth in part by stabilizing *MYB* transcripts.

### Combined editing-expression analyses reveal immune and regulatory pathways altered in B-ALL

Although ADAR1 expression was comparable between B-ALL and T-ALL, B-ALL displayed a lower and less diverse RNA editing landscape compared to T-ALL (Figure 2). When comparing the differential gene expression and differential RNA editing between B-ALL at diagnosis and normal control, a total of 91 genes were both upregulated expression and editing compared to controls (ifdr < 0.05, editing coef. B > 5, exp log fold change > 1), and 38 genes had reduced expression and elevated editing (lfdr < 0.05, editing coef. T > 5, expression log fold change < -1) (Figure S13A). The genes with both elevated expression and editing included B-cell receptor genes and PI3K signaling pathways (*LAT2, PIK3AP1, PIK3CD*). Similar to what was found in TALL, transcriptional and epigenetic regulators including *TMEF2D, CREBBP,* and *MYB* show elevated expression and editing in B-ALL compared to controls (Table S5). While the genes with concurrently elevated expression and editing were significantly interacting in the STRING interaction network (p < 1E-16), indicating the presence of shared underlying pathways (Figure S13B), the downregulated genes were not significantly interacting in STRING (p = 0.2). The genes with concurrently elevated expression and editing were significantly enriched for pathways related to immune system processes (Table S9). The downregulated gene set did not show significant functional pathway or GO term enrichment.

Since A-to-I RNA editing was associated in leukemia stem cell maintenance and disease relapse in several leukemia ^11,23,27,30,52^, we investigated if RNA editing contributes to B-ALL relapse. Paired B-ALL samples at diagnosis and relapse were used to compare the difference in RNA editing landscape. To our surprise, no significant differences were observed in RNA editing percentage at each site (VAF) (Figure S4), overall editing level, number of editing sites, and editing across genic regions in either Alu and non-Alu regions (Figure S14). Notably, there was no significant change in ADAR expression between diagnosis and relapse in B-ALL(Figure 2B), suggesting that RNA editing may not play a major role in B-ALL relapse in contrast to what has been reported in T-ALL^28^.

### Individual RNA editing events associated with patient survival

Overall A-to-I RNA editing and recoding editing are both frequently associated with poor patient survival and inferior clinical outcome in adult malignancies^23, 76, 77^. An important question is whether overall RNA editing, or specific editing events are associated with clinical output in ALL. Here, we leveraged the large sample size of the T-ALL cohort to identify any clinically relevant RNA editing events associated with patient survival. We first integrate the 1,605 differentially edited T-ALL genes (lfdr<0.05, abs (coef. T)>10, Figure S9A) and 581 significantly differentially edited genes in B-ALL (Figure S9C) compared to control (abs (coef. B)>5, FDR<0.05) with survival metadata. The overall editing burden did not predict survival in T-ALL or B-ALL (Figure 7A). Moreover, neither sex nor ethnicity significantly predicted survival in T-ALL or B-ALL (Figure S15).

**Figure 7:**
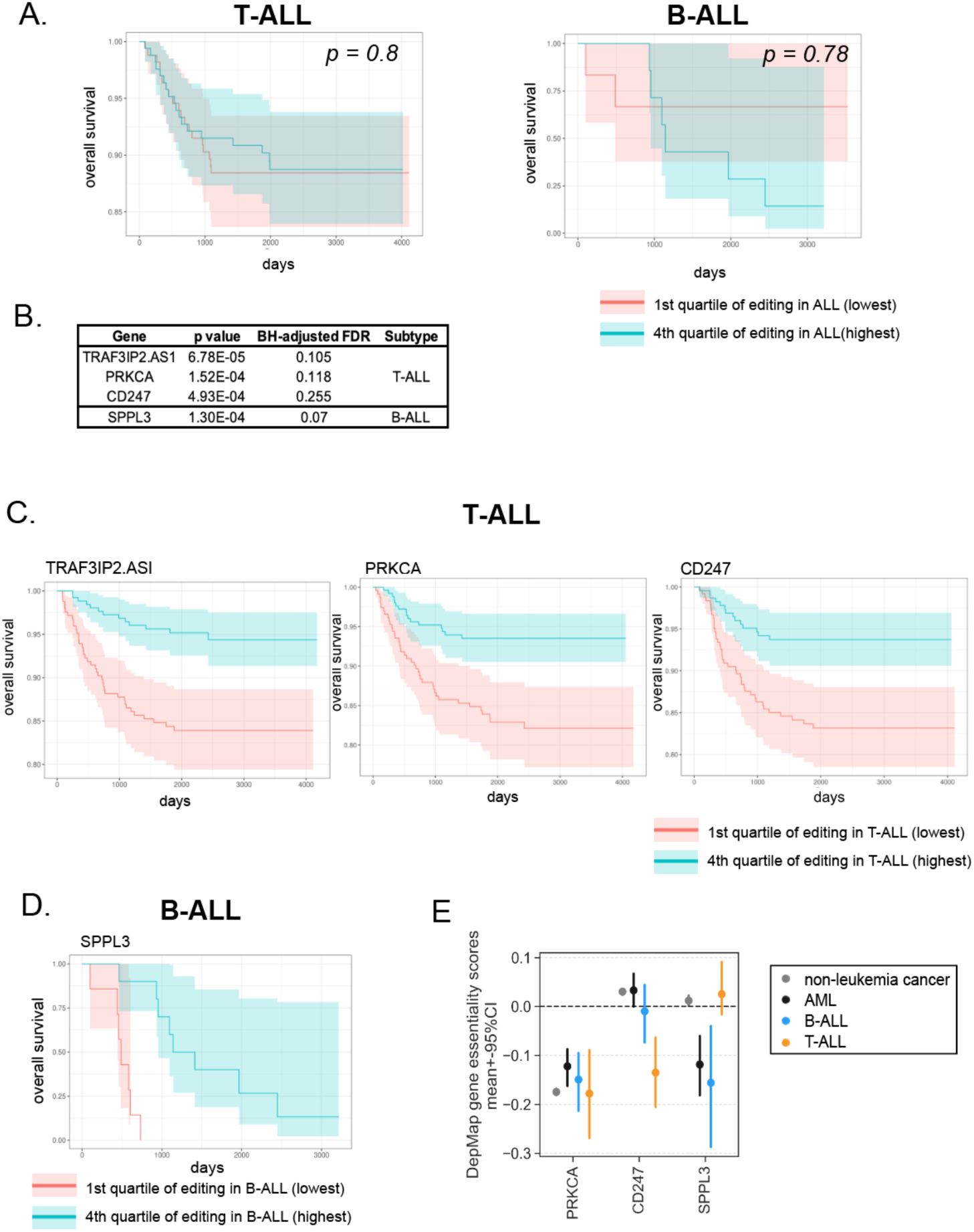
RNA editing in select genes is associated with survival in pediatric ALL. **A.** Kaplan-Meier survival curves comparing patients in the top (4th) and bottom (1st) quartiles of RNA editing levels for overall RNA editing in T-ALL and B-ALL. **B.** Table showing genes with nominally significant associations between RNA editing levels and survival, as determined by Cox proportional hazards regression, including raw p-values and Benjamini-Hochberg adjusted false discovery rates (FDRs). Subtypes refer to the ALL subtype (T-ALL or B-ALL) in which the association was observed. **C-D.** Survival associated with selected genes listed in (B). RNA editing in *TRAF3IP2.ASI, PRKCA*, and *CD247* are associated with better survival in T-ALL (C) and editing in *SPPL3* is associated with better patient survival in B-ALL (D). **E.** DepMap scoring of survival-associated genes in T-ALL, B-ALL, AML, and non-leukemia cancers.

At individual gene levels, we relaxed the FDR cutoff to boost the discovery power. With an FDR of < 0.3, we identify 4 differentially edited genes (*TRAF3IP2.AS1*, *CD247* and *PRKCA* in T-ALL, and *SPPL3* in B-ALL) that were predictive of survival after correcting for multiple testing (Figure 7B-D). These edits were located in intronic and flanking regions of prospective genes (Figure S16). To our surprise, higher editing levels of these four genes were associated with improved survival in T-ALL and B-ALL*. SPPL3 is* a key regulator of cell-surface glycosylation that controls immune recognition and therapy response, particularly in B-cell leukemias^78, 79^. *CD247* (encode T-cell receptor zeta) is a component of the TCR complex and important for T-cell development^58^; *PRKCA* codes for protein Kinase C alpha (PKCa), a serine-threonine kinase that play essential role in single transduction pathway^80, 81, 82^; and *TRAF3IP2* is an adapter protein that mediates cytokine response in response to pathogens and cellular stress^83, 84, 85^. We further examined the essentiality scores of the 4 RNA edited genes in DepMap cancer cell line. *CD247* exhibits increased but moderate essentiality in T-ALL compared to other cancer types, *SPPL3* has specificity in B-ALL and AML, while PRKCA appears to be pan-cancer dependent (Figure 7E). These data suggest that although RNA editing is globally elevated in ALL, specific RNA edited events potentially contribute to favorable patient outcomes.

## Discussion

The A-to-I RNA editing landscape was first systematically profiled across 17 types of adult tumor types using TCGA database^23^, and was later expanded to almost all adult cancer types, including hematological malignancies^86^. Consequently, there is a strong interest in understanding the mechanisms by which ADAR1 influences leukemogenesis. Here, we present the first comprehensive evaluation of A-to-I RNA editing landscape in pediatric leukemia including T-ALL and B-ALL.

Overall, we found that both ADAR1 expression and global RNA editing levels are enriched in TALL relative to normal controls. As previously reported, the majority of the editing (>95%) attributes to Alu regions. Notably, elevated ADAR1 expression coincides with the expression of several critical dsRNA sensors (e.g. PKR and PARP9) and is associated with increased inflammation pathways and interferon signaling. This pattern suggests that ADAR1 represents a cellular response to buffer excessive interferon signaling triggered by endogenous dsRNA in TALL. However, this observation does not explain the heterogeneous RNA editing landscape among T-ALL patients, since neither ADAR1, dsRNA sensors or ISGs were found to correlate strongly with the RNA editing. The discrepancy likely reflects the multi-layer regulation of RNA editing and/or the abundance of dsRNAs in this cellular context. Moreover, patient variability due to different molecular subtypes and treatment-induced stress might further modulate the RNA editing levels. Future studies using single-cell RNA-seq and multi-omics approaches in a large and well-annotated T-ALL cohort with clearly defined treatment history will be crucial to delineate the cellular heterogeneity of ADAR1 activity and dsRNA burden.

In a sharp contrast to T-ALL, B-ALL exhibits similar RNA editing levels and comparable ADAR1 expression to those of normal controls. Unlike hyper-editing in relapsed T-ALL^28^, paired comparison of B-ALL at diagnosis and relapse showed no evidence of increased RNA editing activity during B-ALL relapse. It is worth noting that the number of recoding editing events are comparable between T-ALL and B-ALL on a per sample basis. Rather, the discrepancies are largely driven by Alu-directed RNA editing profiles. It is curious why B-ALL selectively promotes RNA editing in coding regions. These differences may reflect fundamental differences in lineage specificity of leukemia origin and/or molecular mechanisms underlying leukemia relapse. Nevertheless, ADAR1 remains an essential gene in B-ALL according to the DepMap dataset, suggesting it may exert RNA editing-independent functions critical for B-ALL survival. Since RNA edits are often cell-context specific, analyses of lineage-specific RNA editing profiles in normal pediatric hematopoietic populations will help to understand if the observed differences between T-ALL and B-ALL reflect cell type-specific editing preferences or leukemia-associated changes.

RNA recoding edits account for ∼1-2% of all editing events in T-ALL and B-ALL. A previous study examining recoding events reported that recoding in each pediatric cancer subtype resembled those of the normal tissue of origin rather than tumor-specific events^87^. In agreement with our study, many well-known functional recoding events (e.g. *AZIN1, COG3, FLNB*) previously reported were pan-cancer in all pediatric cases^45, 46, 88, 89, 90, 91, 92^. Interestingly, we also found that recoding in T-ALL revealed disease-specific events such as the nonstop edits at *SLFN12L* 3’UTR region. Notably, these edits were also detected in B-ALL but at a much lower frequency. This nonstop RNA editing in translational termination regions could generate longer proteins which would contribute to leukemogenesis. Strikingly, we also reported RNA edits in normal control that are lost during leukemia transformation (e.g. *DACT3*), suggesting RNA editing may also exert protective functions. Whether these edits indicate opportunistic ADAR1 activity based on transcript abundance or are actively targeted in a cell context specific manner will require validation at protein level and follow-up functional studies.

Lastly, we performed survival analysis to assess the clinical relevance of RNA editing events in pediatric ALL. The global RNA editing level was not associated with patient survival. At individual gene levels, higher editing in three genes (*CD247, PRKCA,* and *TRAF3IP2.ASI*) and one gene (*SPPL3)* were associated with improved survival in T-ALL and B-ALL, respectively. These observations suggest that editing at specific genes potentially exerts protective effects against disease progression rather than promoting leukemogenesis..

Together, this study provides the first comprehensive map of A-to-I RNA editing in pediatric leukemia and reveals critical insight into the editing landscape and potential functional targets of ADAR1 in T-ALL and B-ALL. These findings highlight the importance of RNA modification as an additional transcriptional regulation in shaping leukemogenesis and patient outcome.

## Material and Methods

### RNA Editing Detection and Annotation

RNA editing events were identified from paired-end RNA-seq data using a modified SNPiR-based pipeline. FASTQ files were aligned to the human reference genome (GRCh38) with STAR (twopass mode) using GENCODE transcript annotations, followed by duplicate marking with Picard, sorting and indexing with Sambamba, and intron-split adjustment with GATK SplitNCigarReads. Variants were called with GATK HaplotypeCaller and filtered for sequencing artifacts using GATK VariantFiltration (FS > 30.0, QD < 2.0, QUAL < 20, depth < 5 reads). Bases were recalibrated with GATK BaseRecalibrator/BQSR. Candidate editing sites were restricted to known A-to-I editing positions from REDIportal v3, and separated into Alu and non-Alu categories using BEDTools intersections with RepeatMasker annotations.

For Alu sites, SNPiR filtering steps were applied, including removal of mismatches at read ends, exclusion of low-coverage sites (<5 total reads or <2 alternate reads), and filtering of mismapped reads with BLAT. For non-Alu sites, additional filters were applied to remove variants in repetitive elements, intronic sites within 4 bp of splice junctions, homopolymeric regions, and mismapped reads. Final high-confidence Alu and non-Alu editing sites were annotated with Ensembl VEP (cache version 100, GRCh38) and converted to Mutation Annotation Format (MAF) with vcf2maf. A custom R script was used to merge, filter, and produce the final table of RNA editing events for downstream analysis.

### Normalization

To account for variability in sequencing depth across samples, we applied a coverage downsampling procedure to the RNA editing data. For each sample, we first quantified the distribution of site-level read depths (DP) and estimated the 95th percentile coverage (q95). We selected a global target depth of 23 reads, corresponding to the minimum q95 value among samples that could be retained, ensuring that all retained samples could be normalized to the same effective coverage. Samples with q95<23 (n=10) were excluded from downstream analysis, since their coverage was insufficient to confidently detect RNA editing and could not be reliably normalized upward. 1025 T-ALL samples, 74 B-ALL samples (37 diagnosis, 37 relapse), and 6 WT normal were retained for downstream analysis. For each site in each retained sample, we simulated binomial thinning of the read counts, randomly retaining each read with probability 23/q95. This procedure yielded new downsampled total depth (DP) and edited read counts (ALT), bringing sample coverage distributions into close agreement. As illustrated in Supplemental Figure 3, raw coverage distributions (Figure S3A) show substantial systemic differences between samples, which are reduced after downsampling (Figure S3B), demonstrating that the procedure effectively normalizes DP across the cohort. Sites with no downsampled edited reads were discarded. To reduce false positives due to low allelic fractions, we additionally required a variant allele frequency (VAF) >= 0.1, excluding sites with ALT/DP<0.1 after downsampling. The resulting dataset contained coverage-normalized editing calls across T-ALL, B-ALL, and normal samples, suitable for comparisons of overall editing burden, editing burden by edit type (intron, flanking region, missense, splice site, etc), as well as differential editing analysis.

### RNA expression analysis

RNA-seq counts data (T-ALL, B-ALL, WT normals) was downloaded from KidsFirst and Target databases (see data availability). For each sample, we extracted expected counts from RSEM files and assembled them into a combined expression matrix. Gene identifiers were standardized by removing version numbers from Ensembl IDs. The resulting matrices were harmonized across cohorts by intersecting common genes, yielding a unified expression dataset. Expression values were normalized within each cohort using a relative log expression (RLE) procedure: log2transformed counts (log2(counts)+1) were centered by subtracting the per-sample median of high abundance genes (defined by mean log-expression >6-8, depending on group), and rescaled to produce normalized expression matrices. Differential expression analysis was then performed between groups (e.g. T-ALL vs normal, B-ALL vs normal), using linear modeling (limma^93^). To correct for multiple testing, we estimated local false discovery rates (lfdr) using the q value package in R, and reported lfdr-adjusted significance values alongside raw p-values.

### Differential editing analysis

#### Gene-level differential editing (coding and noncoding)

To assess differential RNA editing between groups, we constructed a gene-by-sample editing matrix from the downsampled MAF files. For each sample, edited sites were collapsed by genes, and the number of edited sites per gene was recorded. These counts were assembled into a gene x sample matrix, and aligned to the corresponding normalized expression matrix. Genes with poor coverage were excluded prior to modeling. Specifically, genes with > ⅔ missing values in either the case or control group were removed, as these lacked sufficient data to support reliable testing. To test for differences in editing burden between conditions (e.g. B-ALL vs normals, T-ALL vs normals, B-ALL relapse vs diagnosis), we fit linear models of the form *editing ∼ condition + expression*, where editing is the per-gene editing count and expression is the normalized expression level of the same gene. This approach accounts for gene expression level as a covariate, to avoid spurious associations arising from expression-dependent detection of edits. Regression coefficients (coef.T, coef.B, etc) for condition terms represent the direction and magnitude of differential editing, and p-values for these coefficients were obtained directly from the linear models. To correct for multiple testing, local false discovery rates (lfdr) were estimated using the qvalue package, and lfdr-adjusted significance values were reported alongside raw pvalues.

#### Note on imbalanced statistical testing

When comparing T-ALL and B-ALL samples to normals, there is a large imbalance in sample sizes. To mitigate bias due to this imbalance, we adopt the procedure recommended by Efron and Tibshirani in chapter 16 of their book on bootstrap ^51^. Briefly, we first calculate the coefficient representing *condition* in the linear model *editing ∼ condition + expression*, using the available data, for every gene. Specifically, in the comparison of T-ALL to controls, we set *condition=0* for controls and *condition=1* for T-ALL. The regression coefficient associated with *condition* we call *coef.T*. We store *coef.T* for later. Next, we adjust control samples by adding *coef.T* to the *editing* variable in all control samples. We create a null ensemble by simultaneously bootstrapping the adjusted controls and cases. The null *coef.T* is calculated and its distribution is stored for every gene. We performed 10,000 bootstrap samples for both T-ALL- and B-ALL-vs-control comparisons. We calculate the marginal *p*-values as the fraction of null *coef.T* values at least as extreme as the observed *coef.T*. In this case, the effect is so large that the observed value of *coef.T* is outside the range of observed null values for most genes, and the marginal *p*-values are zero (Figure S5). The apparently uniform tails of the distributions of marginal *p>0* allow us to estimate the null gene fractions π_0_ to be ∼0.059 for T-ALL vs. controls and ∼0.31 for B-ALL vs. controls. We note that these are much smaller that the customary situation π_0_∼ 1. The false discovery rate (FDR) at any level of *p* can be estimated as the ratio of expected null tests with *p*value less than *p* (which equals π_0_*p*), and the observed number of tests with *p*-value less than *p*. We also calculate the *z*-score per gene as *z = coef.T/sd(null coef.T)*.

#### Differential editing in coding regions

To specifically evaluate editing differences in coding regions, we filtered the combined edits table (T-ALL, B-ALL, and normal samples) to retain only protein-coding variant classes (Missense, silent, splice region, splice site, and nonstop edits), excluding intronic, UTR, and flanking region edits. Because coding edits are rare compared to noncoding edits, per-gene regression models often resulted in sparse matrices with many missing values, limiting power to detect differences. To address this, we adopted a contingency table framework that operates at the level of gene-edit type-Alu status combinations. Editing events were grouped by a composite identifier combining gene symbol, edit classification, and Alu status (Alu vs non-Alu). For each group, we counted the number of edited sites observed per condition, as well as the total number of possible sites (defined as the number of unique editing sites in the tested region, across both conditions, multiplied by the number of samples in that condition). We then tested for differential editing frequencies between conditions (T-ALL vs normal, B-ALL at diagnosis vs normal, and T-ALL vs B-ALL) using chi-square contingency tests as previously published^26^. To ensure robustness, groups with only a single observed edit were excluded from testing. Multiple hypothesis correction was performed using the Benjamini-Hochberg false discovery rate, and adjusted p-values were reported alongside raw p-values and chi-square statistics.

### DepMap analysis

We leveraged publicly available CRIPSR-Cas9 gene essentiality data from the DepMap project (DepMap Public 23Qx release) to evaluate whether candidate genes identified in our differential RNA editing and expression analyses are required for leukemia cell growth. Gene effect scores (CERES dependency scores) were extracted for the focal gene set across all available cell lines and linked to cell line annotations, including lineage, primary disease, and subtype classifications. Scores were grouped by leukemia subtype, and mean essentiality values with 95% confidence intervals were visualized with a reference line at 0 (0 score meaning the gene was not essential). Gene essentiality scores which were more negative indicated greater essentiality, with a threshold of -1 corresponding to complete loss of viability upon knockout. These analyses provided orthogonal evidence for the relevance of candidate genes in T-ALL biology by integrating functional screening data with editing and expression signatures.

### Survival analysis

RNA editing data was integrated with clinical metadata from T-ALL and B-ALL patients into a survival analysis framework. For each patient, overall survival time was defined as the number of days from diagnosis to last known follow-up or death. Survival was tested for association with zscore of top differentially edited regions, total editing burden, or gene-level editing counts, which were further stratified into quartiles. Individual genes were tested which had significantly changed editing levels between T-ALL and normal (abs(coef.T)>10 and lfdr<0.05) and between B-ALL and normal (abs(coef.T)>3 and lfdr<0.05). Kaplan-Meier survival curves were generated to compare groups with higher versus lower editing levels. In addition, Cox proportional hazards regression models were fitted to estimate hazard ratios and 95% confidence intervals for editing-associated predictors, adjusting for age and sex.

### Pathway/network analysis

Genes sets of interest were subjected to pathway and network analysis using the STRING webtool^94^ (v12). Lists of genes identified from differential expression and differential editing were input to the webtool, using species Homo Sapiens. Network interactions were considered if they had scores of 0.4 or greater (medium confidence). Significance of numbers of interactions were assessed by STRING (permutation test). Pathway analysis was performed by STRING, using over-representation analysis, querying against known pathway databases (including GO, REACTOME, KEGG, wikipathways, pubmed).

### Lentiviral transduction and Western Blotting

SUP-T1, Jurkat, and CUTLL1 T-ALL cell lines were purchased from ATCC and cultured in RPMI 1640 medium supplemented with 10% fetal bovine serum (FBS) and 1% penicillin–streptomycin, and maintained at 37 °C in a humidified incubator with 5% CO₂. Cells (1×10^6^) were transduced with lentivirus at a MOI of 20 in 96-well U-bottom plates in 100 µL culture media for 48 hours. Cells were lysed in RIPA buffer (50 mM Tris-HCl pH 7.4, 150 mM NaCl, 1% NP40, 0.5% sodium deoxycholate, 0.1% SDS) supplemented with protease and phosphatase inhibitors. The lysates were centrifuged at 10,000 × g for 10 min at 4 °C, and the supernatant was collected. The protein concentrations were determined using a BCA assay. Lysate (10 µg) was separated on 10% SDSPAGE gels and transferred to PVDF membranes (Millipore), followed by blocking with 5% BSA in TBST (20 mM Tris-HCl, 150 mM NaCl, 0.1% Tween-20, pH 7.6) for 30 minutes at room temperature. Primary antibodies (MYB, Thermo Fisher, MA1-90154, 1:500 dilution; beta-actin, Cell Signaling, #4976, 1:2000 dilution) were added and incubated with the membrane overnight at 4 °C and then HRP-conjugated secondary antibodies (1:5000, Cell Signaling) for 1 hour at room temperature. Blots were visualized on a ChemiDoc system (Bio-Rad).

## Data availability

The normal pediatric PBMC and B-ALL RNA-seq datasets were obtained from NCI TARGET. The results published here are in whole or part based upon data generated by the Therapeutically Applicable Research to Generate Effective Treatments (https://www.cancer.gov/ccg/research/genome-sequencing/target) initiative, phs000464 and phs000471. The data used for this analysis are available at the Genomic Data Commons (https://portal.gdc.cancer.gov). The T-ALL RNA-seq dataset was obtained and analyzed based in whole or in part upon data generated by Gabriella Miller Kids First Pediatric Research Program projects phs002276, and were accessed from the Kids First Data Resource Portal ( https://kidsfirstdrc.org/ and/or dbGaP (www.ncbi.nlm.nih.gov/gap). The T-ALL with ADAR1 knockdown was obtained from GEO:GSE221112.

## Acknowledgement

This work was supported by Leukemia Research Foundation (Q.J.), Hartwell Foundation (Q.J.), American Society of Hematology (Q.J.), CureBound Discovery Grant (Q.J.) NIH NCI 1R01CA282792-01A1 (Q.J.) and NIH NCI 1R03CA287274-01 (Q.J.), the Swedish Childhood Cancer Foundation/Barncancerfonde (F.H: TJ2014-0014, PR2017-0086), Åke Wiberg Foundation (M18-0151) and Märta och Gunnar V. Philipson Foundation (A.N, F.H). The project was partially supported by the National Institutes of Health, Grant UM1TR005449 and UM1TR005449. The content is solely the responsibility of the authors and does not necessarily represent the official views of the NIH.

## Author Contributions

S.B.R., T.M., R.S.., M.R., G.I.D., C.R. and S.E. performed the computational analysis and experiments. S.B.R., D.K., F.H. and Q.J. conceptualized the project and wrote the paper. D.K. and D.T. contributed with scientific expertise and were involved in writing and reviewing the paper.

## SUPPLEMENTAL INFORMATION

### Supplemental Table

Table S1: Patient Demographic summary and normalized edits in Alu and non-Alu regions.

Table S2: Gene expression and pathway (R > 0.25 and p adj < 0.25) associated with ADAR1 expression in T-ALL.

Table S3: Gene set and pathway associated with overall RNA editing in Alu and non-Alu sites. The pathways were analyzed based on top 500 positively and negatively associated genes.

Table S4: Recoding RNA editing events.

Table S5: Differential editing and differentially expressed genes in ALL vs normal.

Table S6: DepMap essential genes selected by elevated editing and expression in T-ALL (coef. T > 10, log FC > 1, lfdr < 0.05)

Table S7: Pathways enriched in differentially edited and differentially expressed targets between T-ALL and normal control.

Table S8: Pathway enriched in overlapping genes between T-ALL vs normal and T-ALL after ADAR1 knockdown.

Table S9: Pathways enriched in B-ALL compared to normal control.

### Supplemental Figure

**Supplemental Figure 1.**
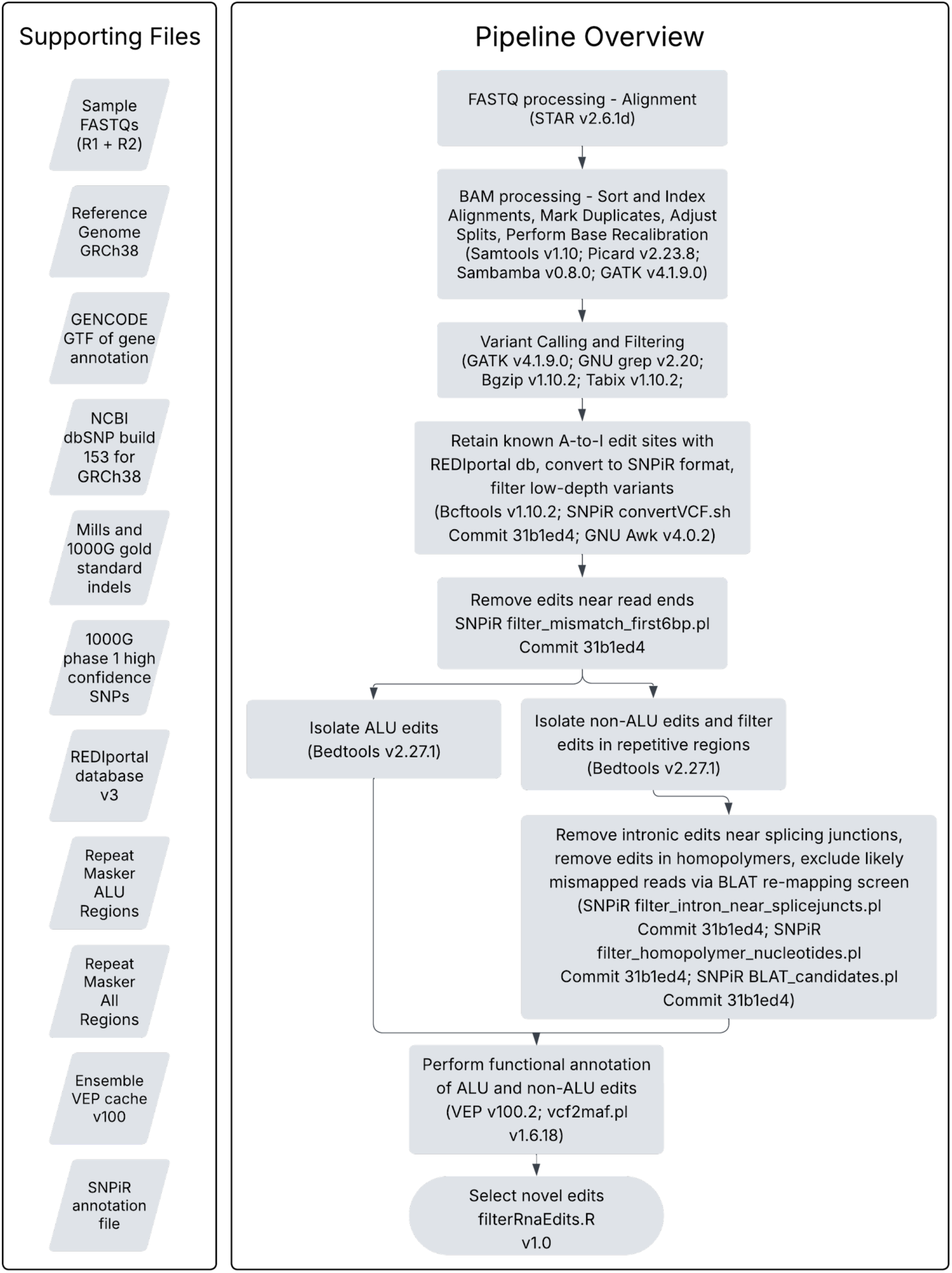
Development of RNA editing pipeline.

**Supplemental Figure 2:**
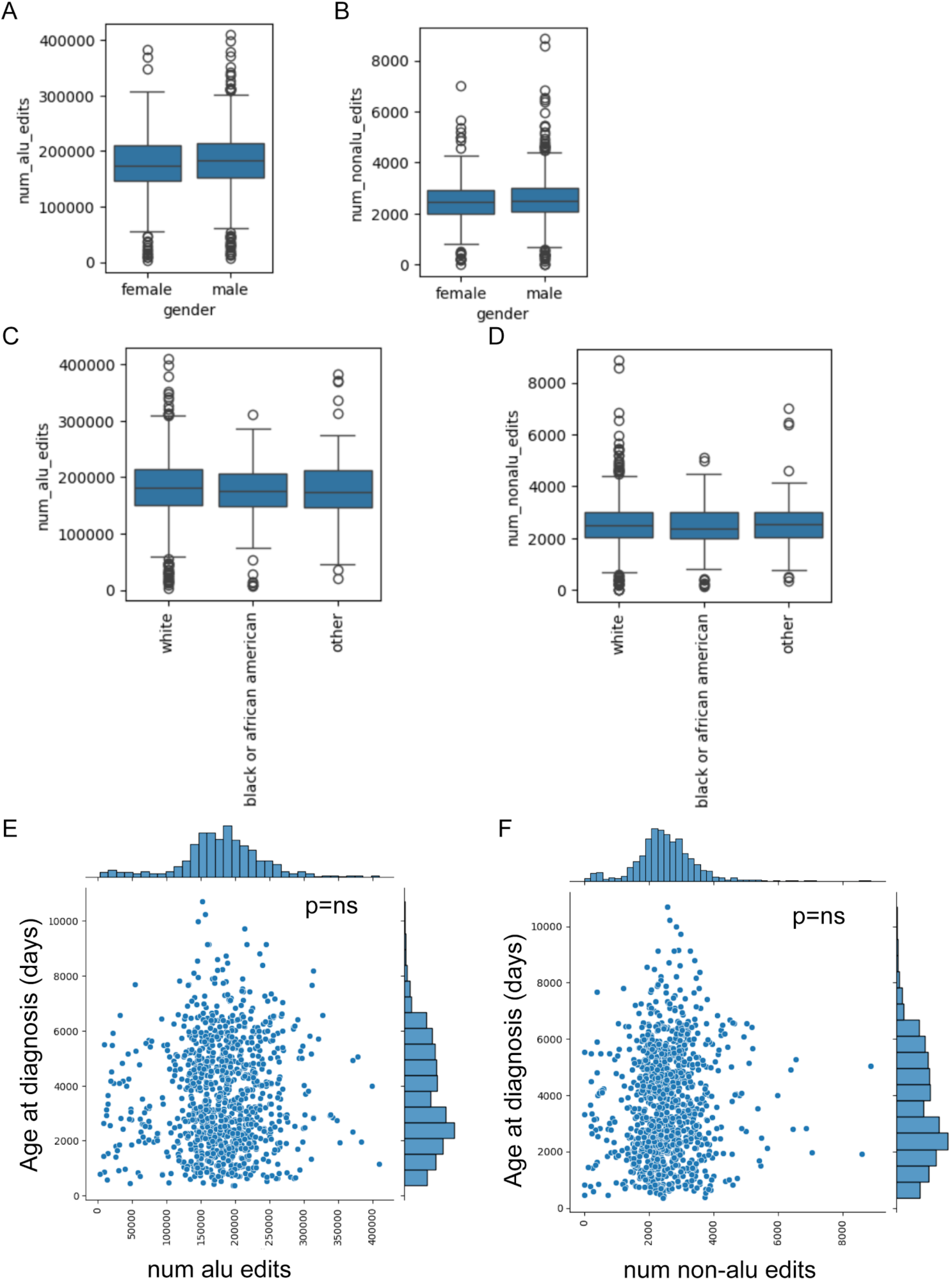
Overall RNA editing landscapes based on clinical information. RNA editing profiles stratified based on sex (A-B), ethnicity (C-D), and age (E-F). Editing in Alu and non-Alu regions are shown separately.

**Supplemental Figure 3.**
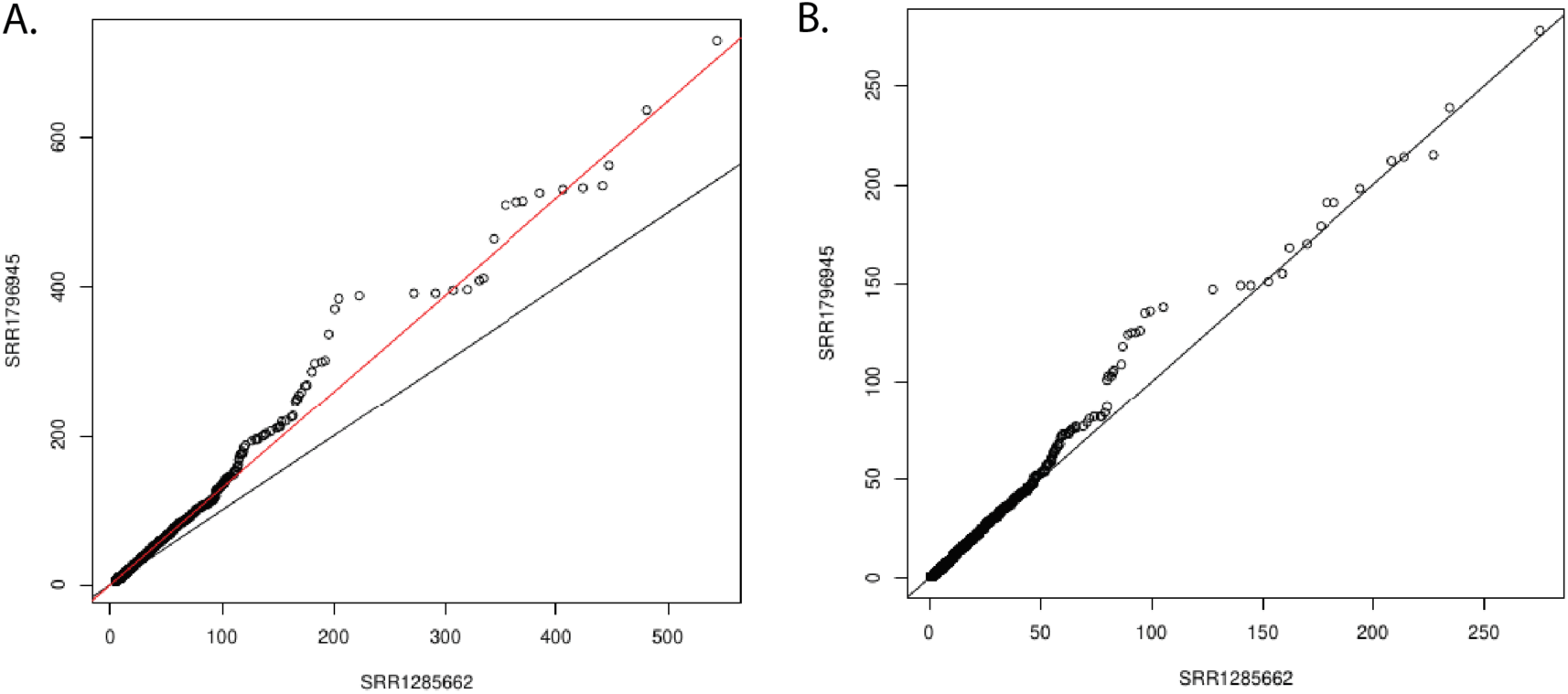
Q-Q plot of DP between two samples for normalization. **A.** Raw data with he black line has a unit slope and the red line has a slope given by the ratio of 95-percentiles of DP in both datasets. **B.** Normalization by downsampling using the probability calculated from the ratio of 95-percentiles of DP distributions.

**Supplemental Figure 4.**
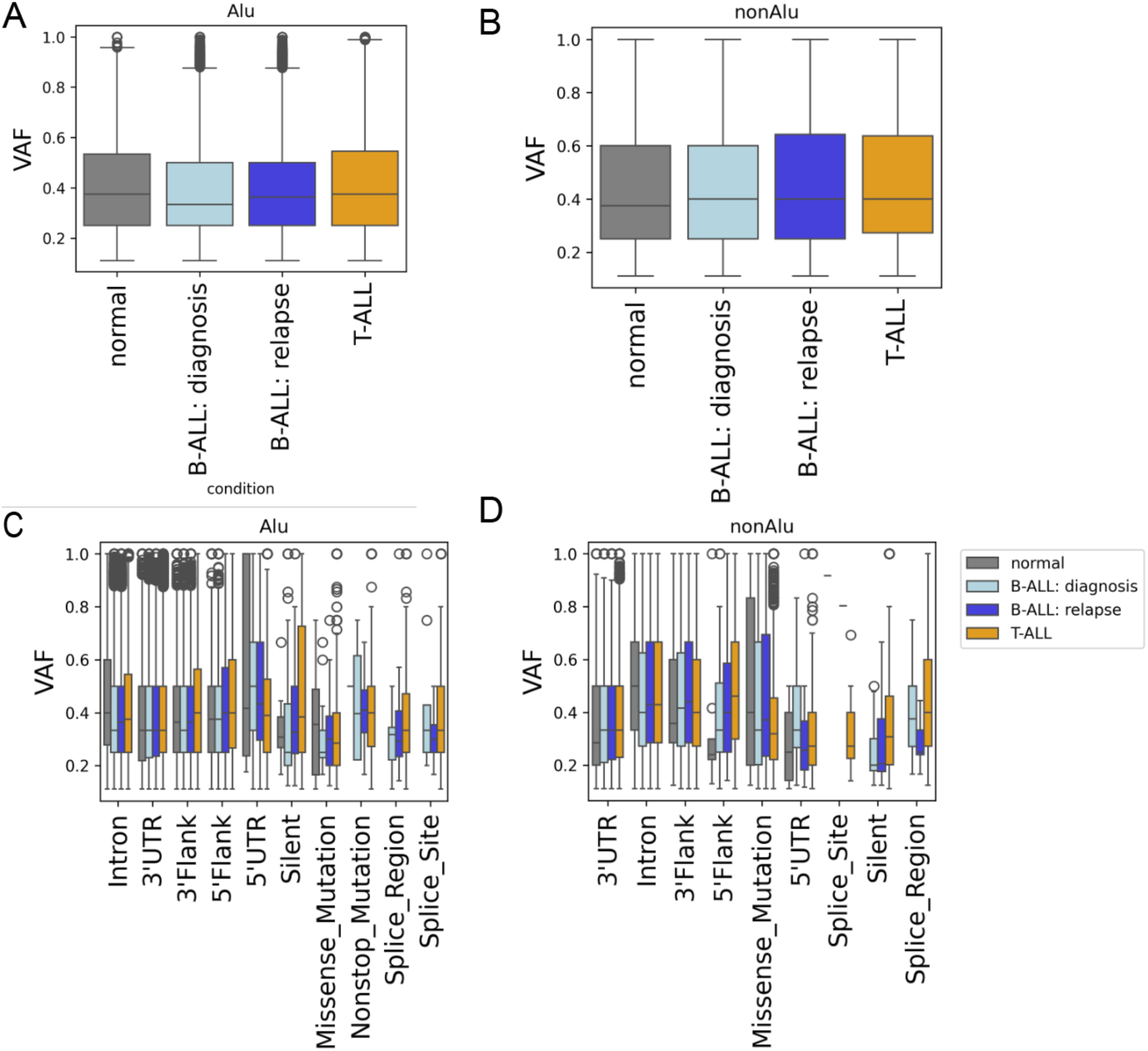
RNA editing level of Variant Allele Frequency (VAF) in nonleukemic normal, diagnosis and relapsed B-ALL, and T-ALL. **A-B.** RNA editing frequency (VAF) in different disease cohorts (n = 6 health control, n = 37 matched diagnosis and relapsed B-ALL, and n = 1,025 T-ALL) in Alu (A) and non-Alu locations (B). **C-D.** Boxplot showing the VAF at various transcript locations of Alu (C) and non-Alu regions (D).

**Supplemental Figure 5.**
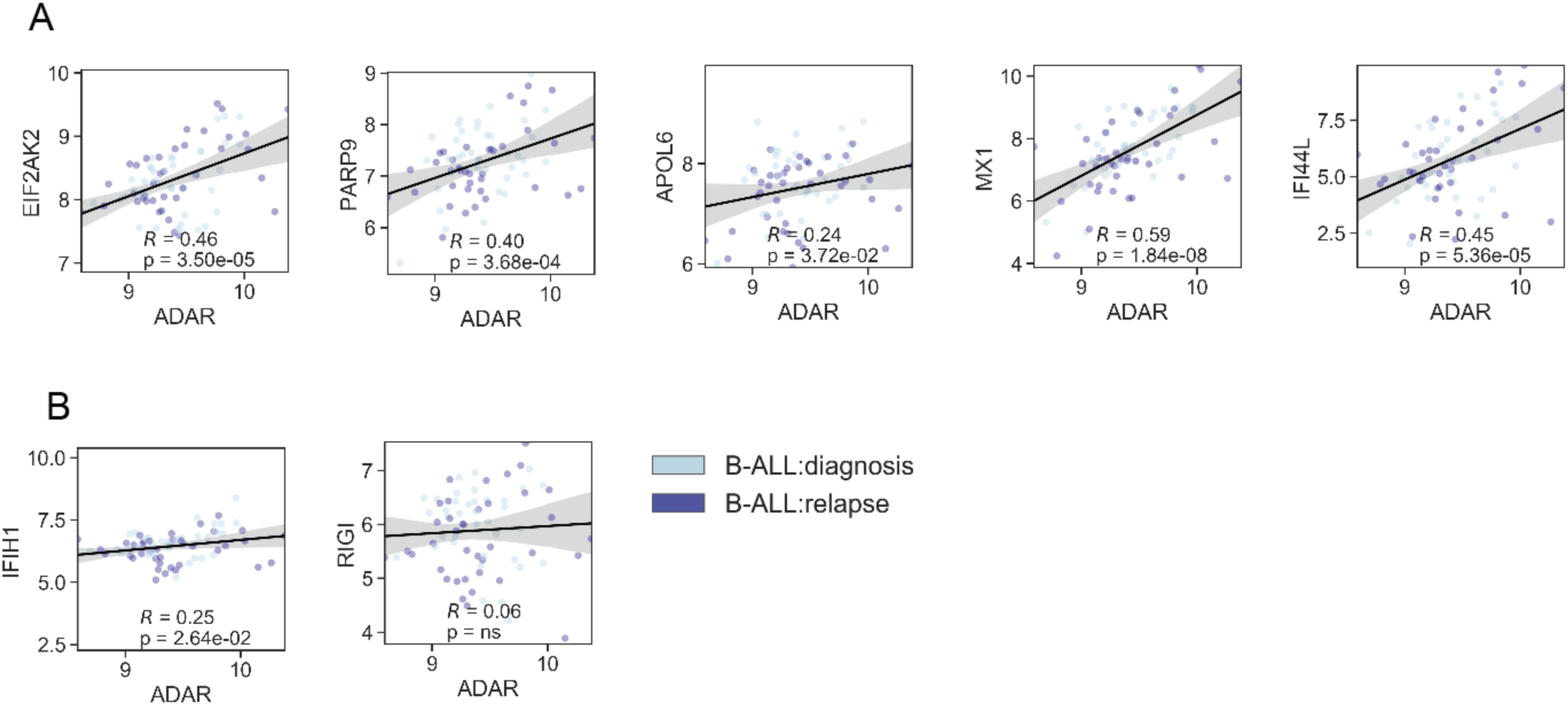
Gene correlation with ADAR1 expression in B-ALL. Top associated genes (A) and known dsRNA sensors (B) were shown. Diagnosis and relapsed cohorts are colored as light blue and purple, respectively.

**Supplemental Figure 6.**
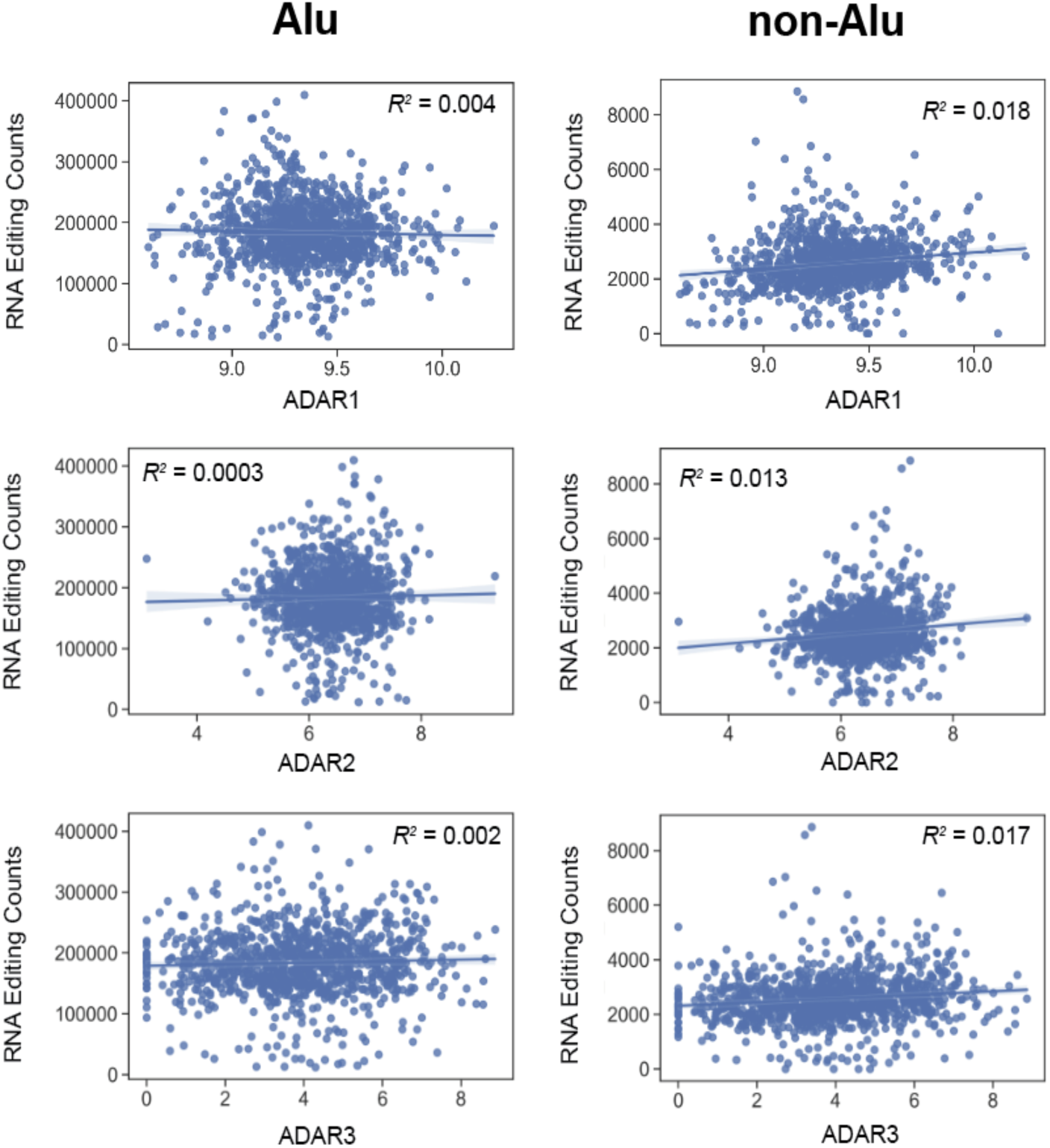
Correlation of gene expression with overall RNA editing. Linear regression model was applied to quantify gene correlation with RNA editing counts in Alu and non-Alu in T-ALL samples.

**Supplemental Figure 7.**
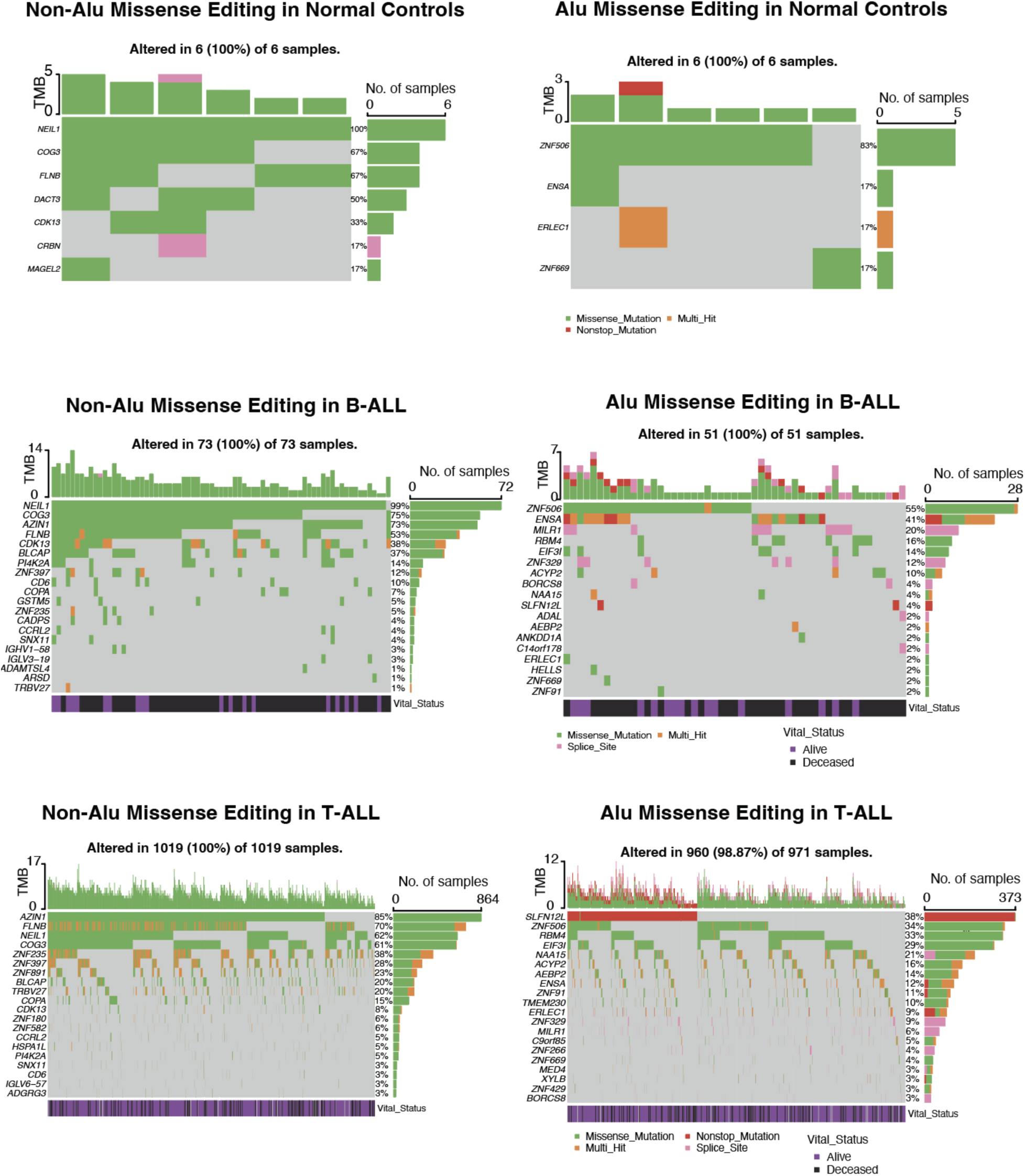
RNA recoding events in normal control and ALL. The editing events were separately shown in Alu and Non-Alu regions.

**Supplemental Figure 8.**
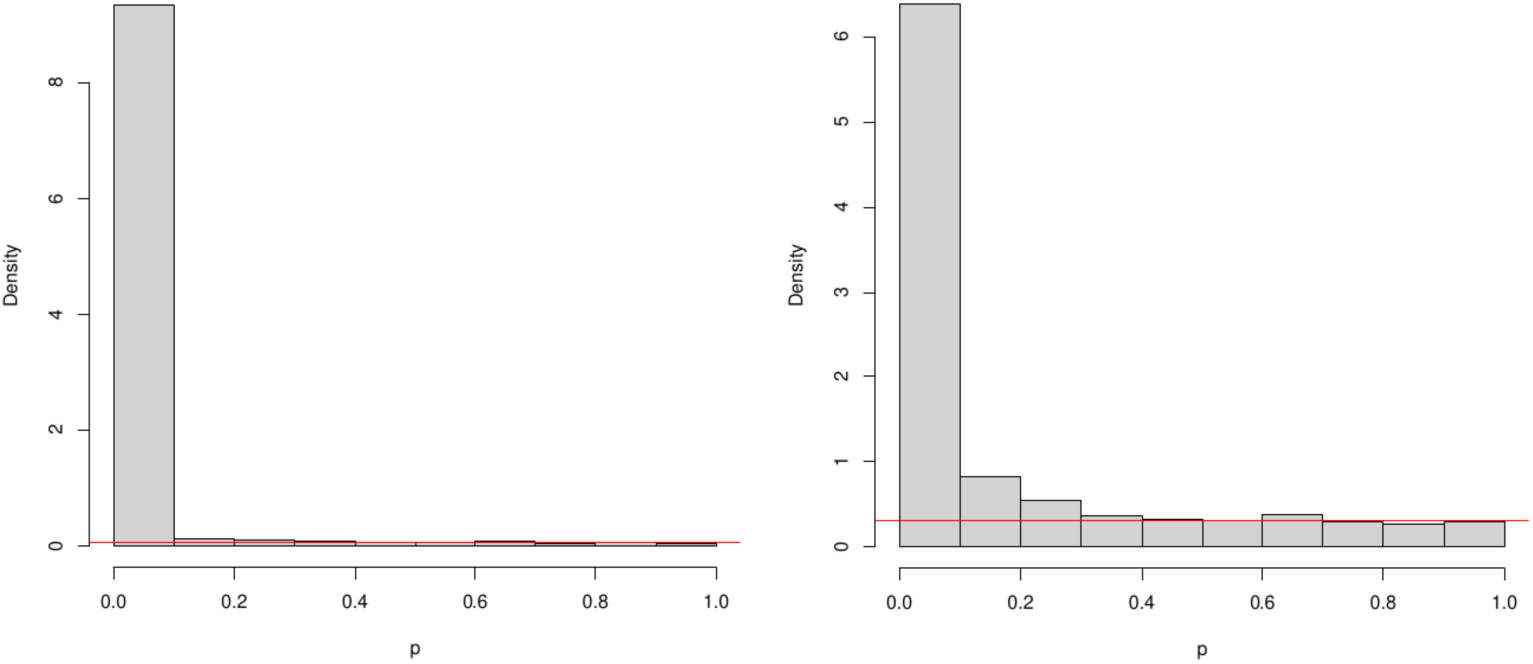
Normalization to correct for sample size imbalance. Distribution of marginal p-values in the T-ALL vs. controls (left panel) and B-ALL vs. controls (right panel) after applying null-centered stratified bootstrap framework. The red lines are drawn at levels 0.059 and 0.31, respectively.

**Supplemental Figure 9.**
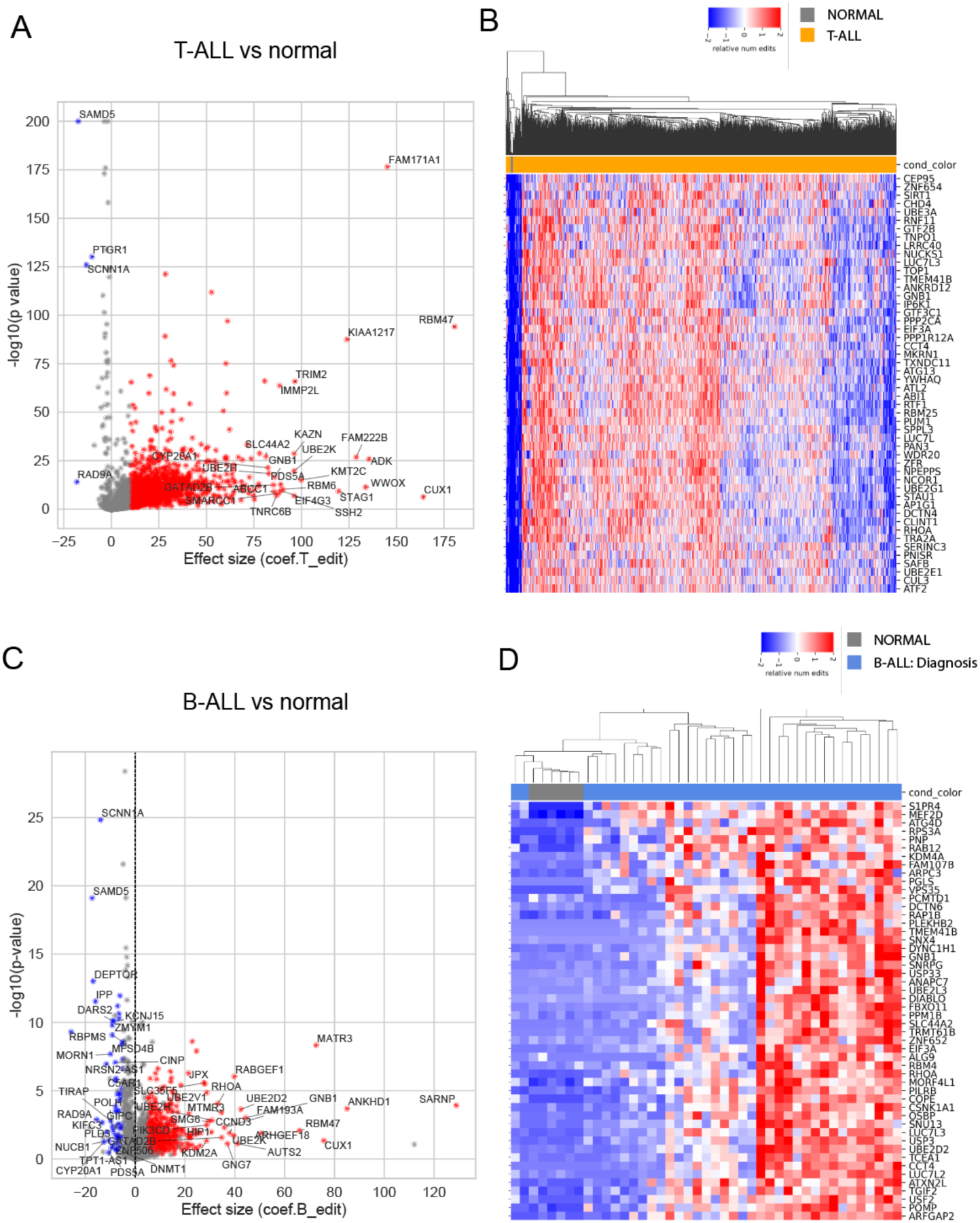
Differentially edited transcripts between ALL and normal control. **A.** Volcano plot of T-ALL vs normal differential editing. Red points indicate coef.T>10 and lfdr<0.05. Blue points indicate coef.T<-10 and lfdr<0.05. **B.** Heatmap showing the relative editing burden of the top 50 genes with most elevated editing levels in T-ALL compared to controls (coding + noncoding). Columns represent samples, and are annotated by condition. Rows represent genes. Both rows and columns were ordered by hierarchical clustering. Heatmap values are z-scores of normalized editing levels, calculated across each row. **C.** Volcano plot of B-ALL diagnosis vs normal differential editing. Red points indicate coef.T>5 and lfdr<0.05. Blue points indicate coef.T<-5 and lfdr<0. **D.** Heatmap showing the relative editing burden of the top 50 genes which had the most elevated editing levels in B-ALL at diagnosis comparing to controls (coding + noncoding). Columns represent samples, and are annotated by condition. Rows represent genes. Both rows and columns were ordered by hierarchical clustering. Heatmap values are z-scores of normalized editing levels, calculated across each row.

**Supplemental Figure 10:**
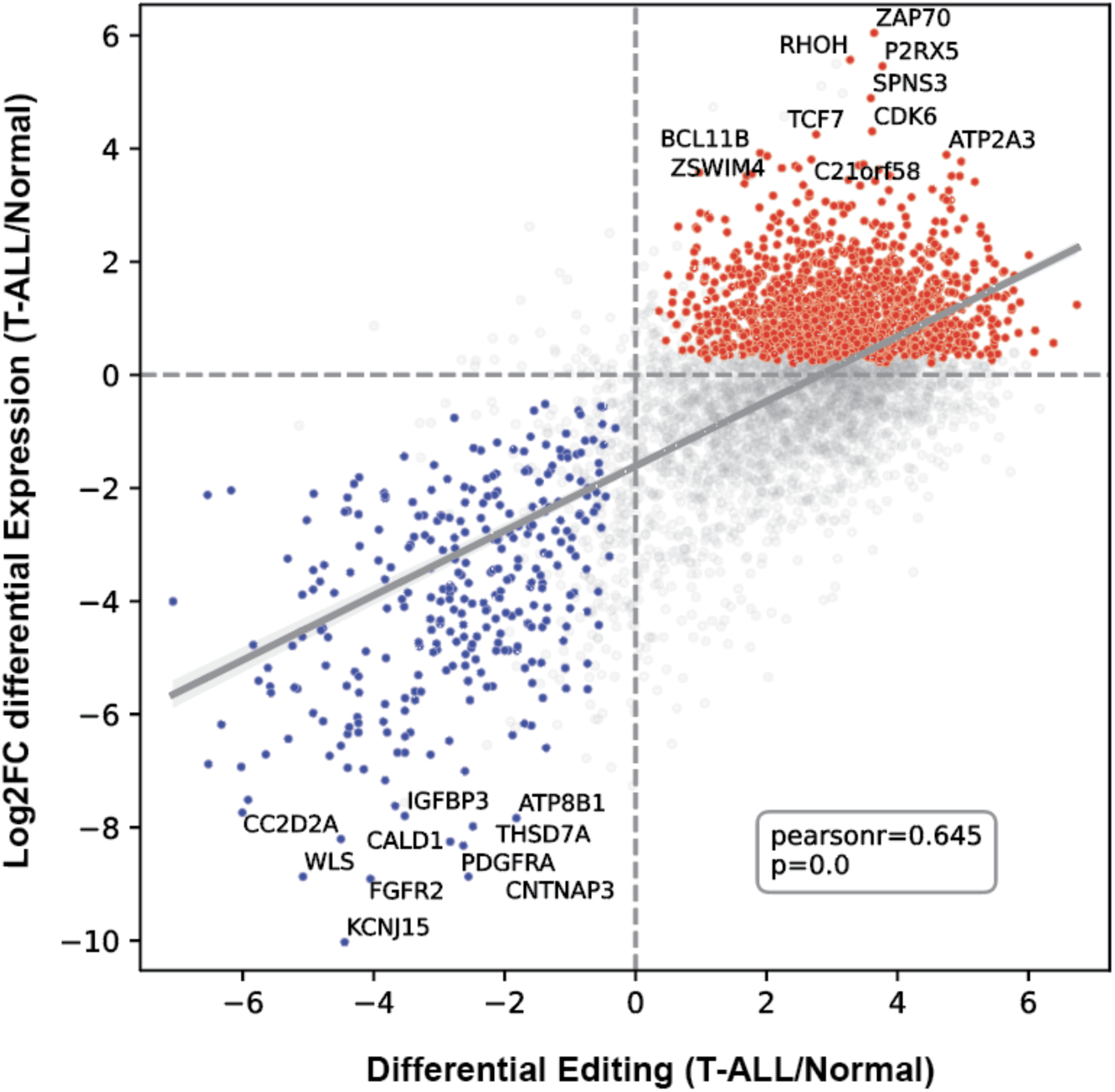
Explore differential A-to-I RNA editing and differential gene expression in T-ALL. The differential expression and the differential edits in each gene was correlated with between T-ALL and normal control.

**Supplemental Figure 11.**
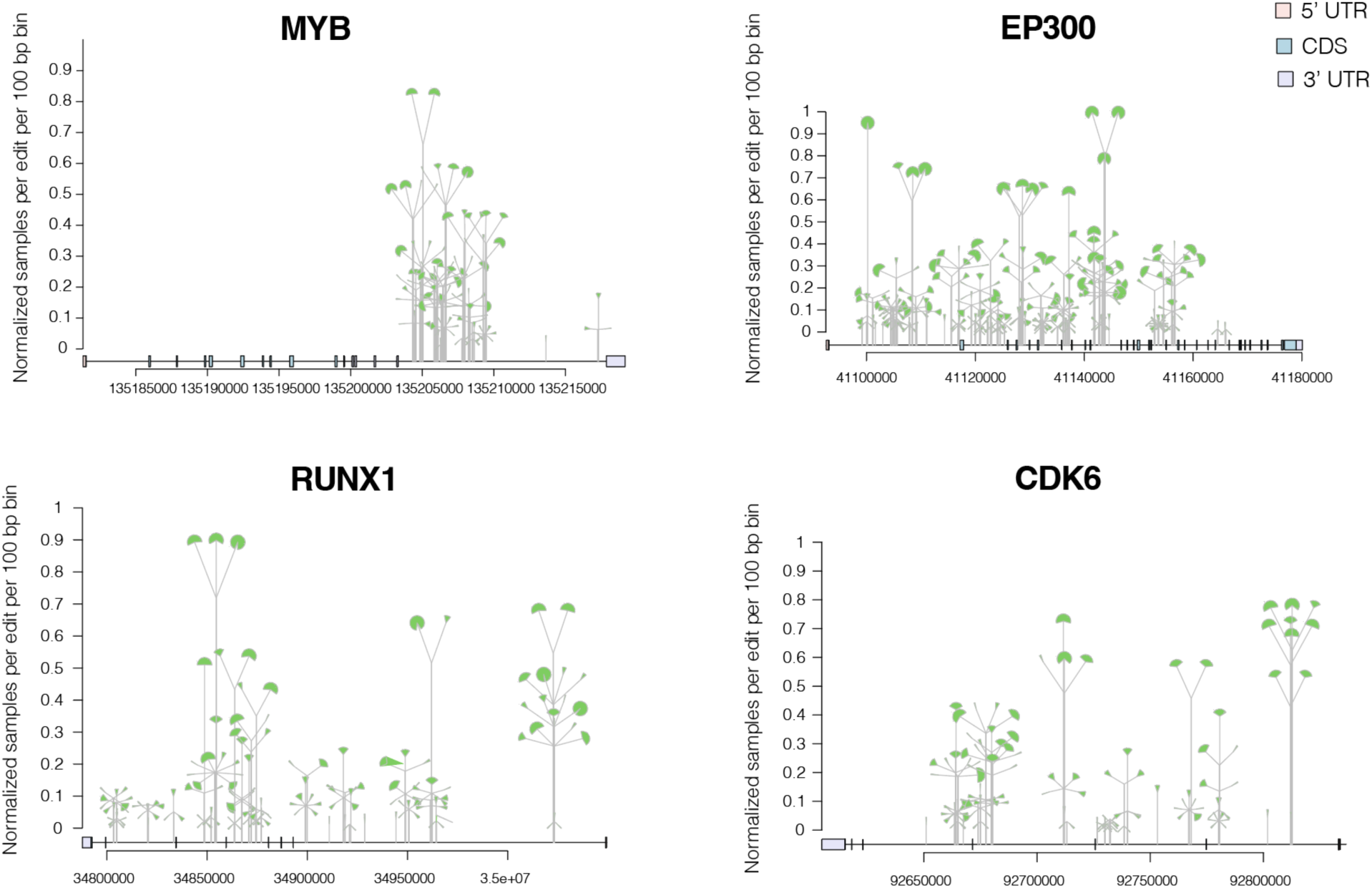
Dedalian plot of editing location with transcripts that are highly differentially edited and expression between T-ALL and normal controls. At the top of each stem, a pie chart shows the fraction of T-ALL or normal individuals that have an edit at the site. The y-axis denotes the normalized edit prevalence (fraction of samples edited) for each genomic bin. Gene models are shown below each plot.

**Supplemental Figure 12.**
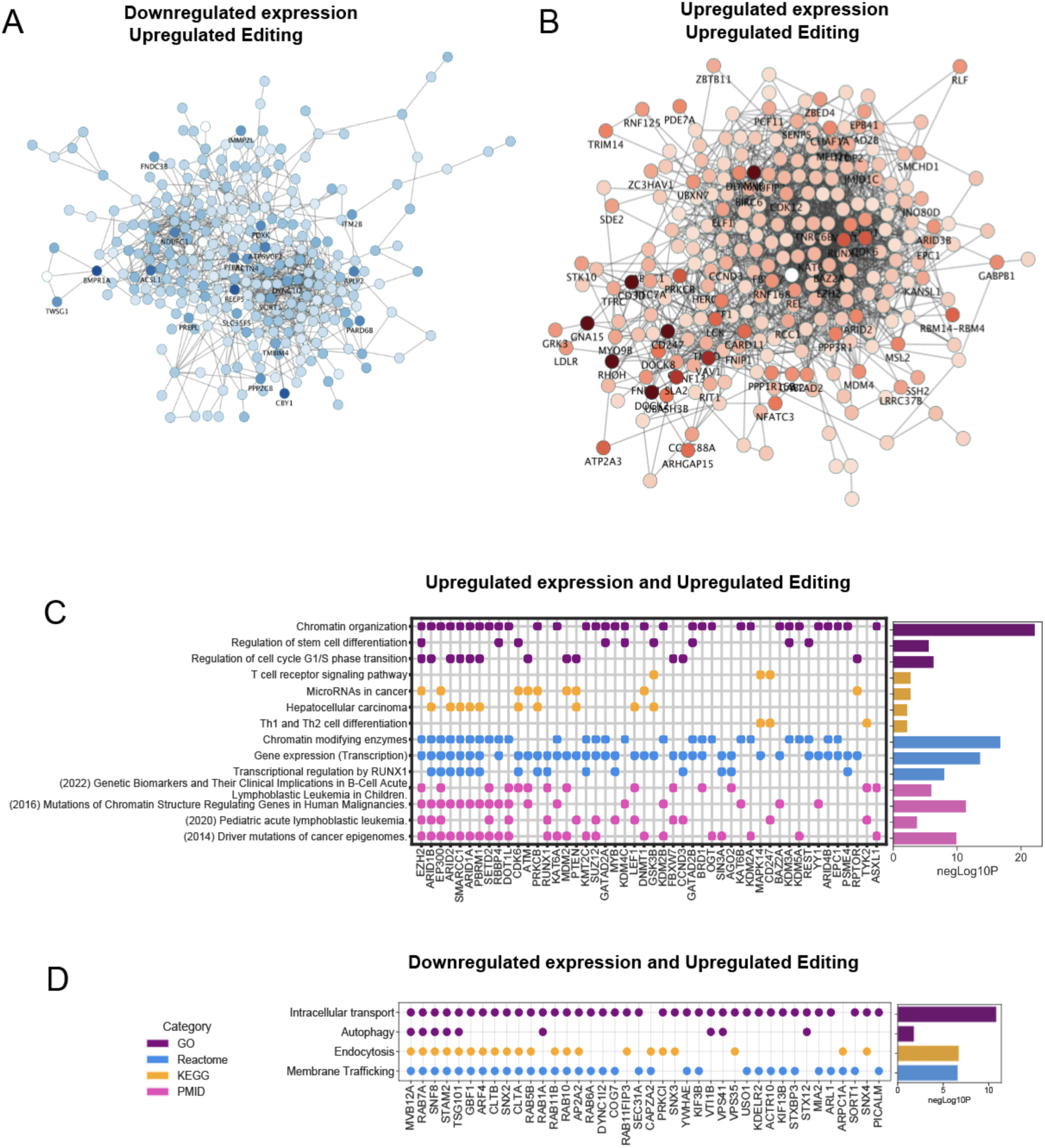
STRING network analysis of differentially edited and expressed genes between T-ALL and normal control. **A-B.** Molecular interaction network (STRING high confidence) of genes with elevated editing in T-ALL compared to normal and A) downregulated expression, or B) upregulated expression. **C.** Pathway enrichment of genes with increased editing and increased expression (red dots in main Figure 5A). Circle colors represent source database; bar plots to the right indicate significance of pathways. Genes are shown if they are present in 3 or more pathways shown. Circle colors represent source database; bar plots to the right indicate significance of pathways. Genes are shown if they are present in 4 or more pathways shown. **D.** Pathway enrichment of genes with increased editing and decreased expression (blue dots in main Figure 5A).

**Supplemental Figure 13:**
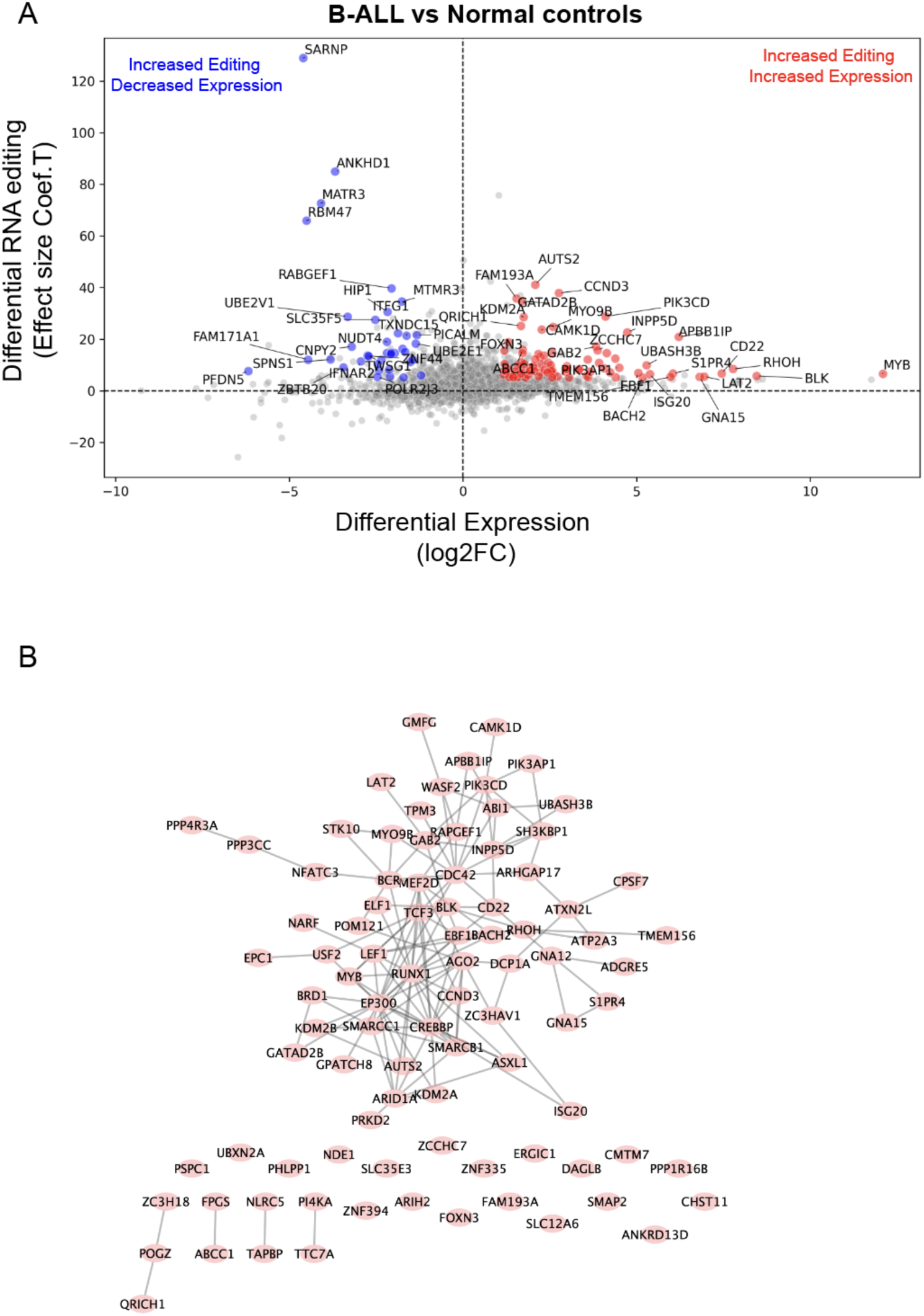
Differentially expressed and differentially edited genes between B-ALL and normal controls. **A.** Scatterplot showing a positive correlation between differentially expressed genes (x-axis, log2 fold change) and differentially edited genes (y-axis, coef. B). Red dots indicate concurrently significantly upregulated genes and significantly more edited genes (FDR <0.05, editing coef. B > 5, logFC > 1). Blue dots show significantly downregulated genes and more edited genes (FDR <0.05, coef. B > 5, logFC < -1). Select genes are labeled. **B.** STRING network analysis of differentially edited and upregulated expressed genes between B-ALL and normal control.

**Supplemental Figure 14:**
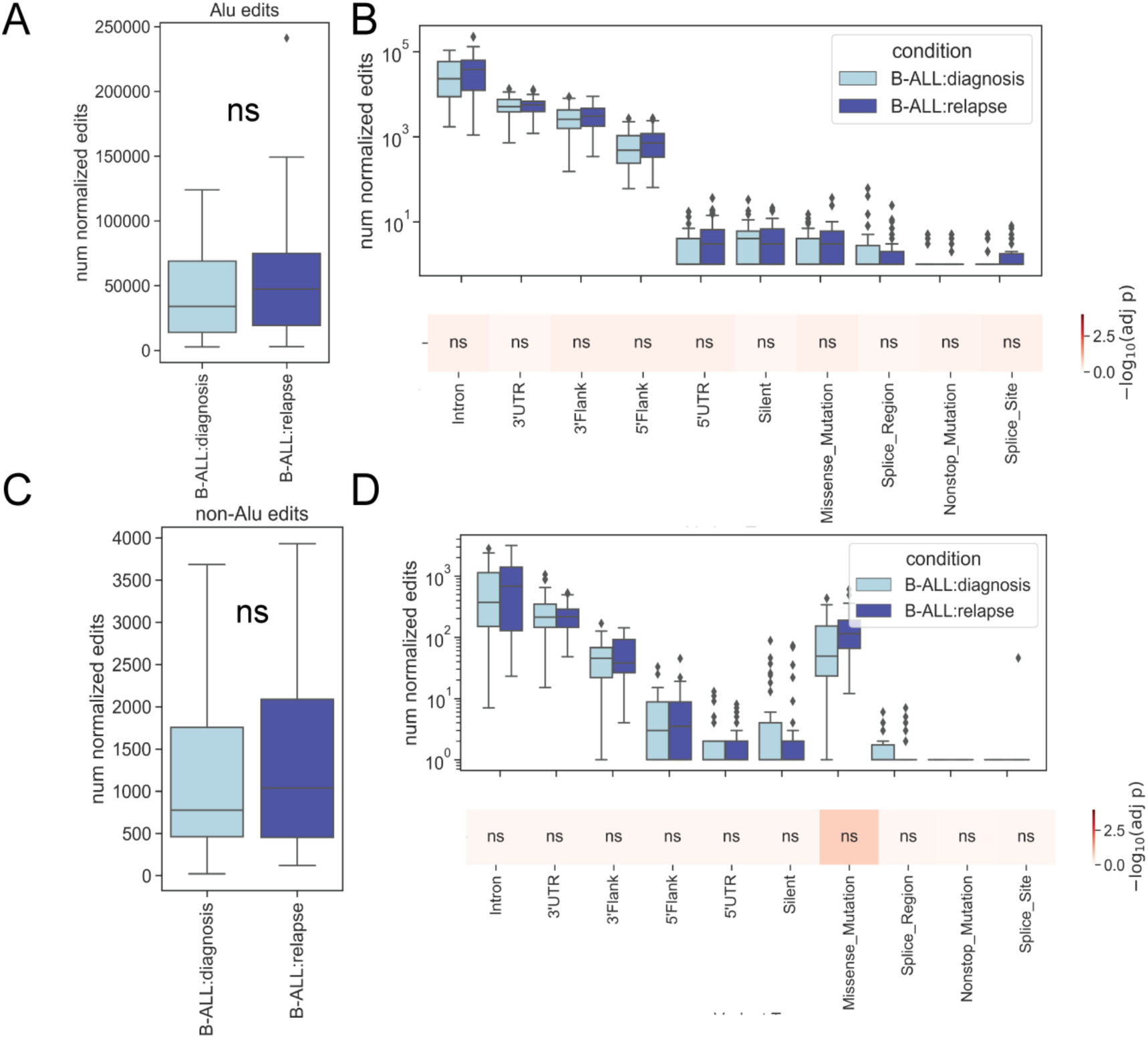
Editing levels between B-ALL diagnosis and relapse. No significant differences were detected across Alu and non-Alu editing events.

**Supplemental Figure 15:**
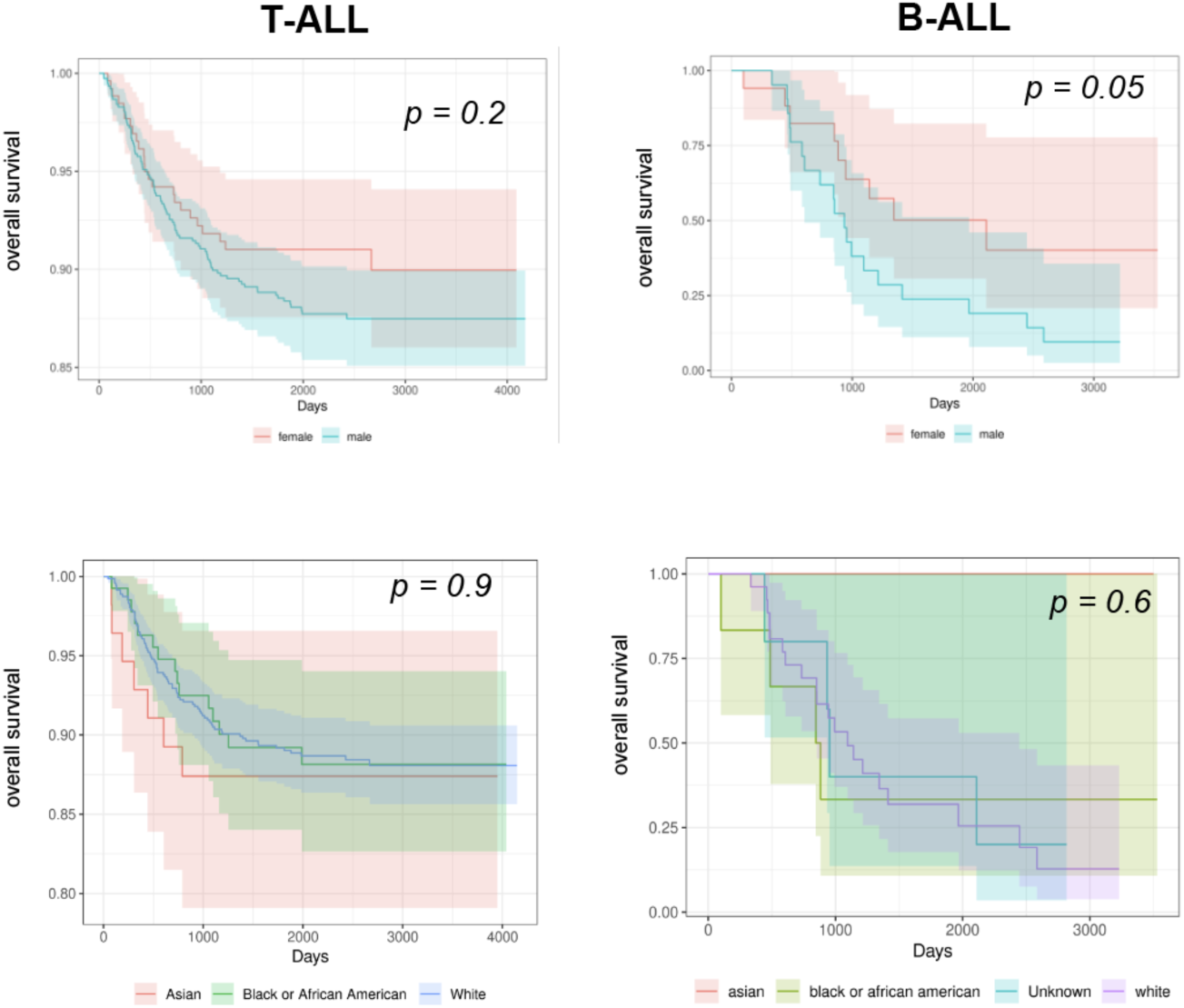
RNA editing levels stratified by sex or race are not associated with survival. The RNA editing level in different sex and ethnicity groups were integrated with clinical metadata in T-ALL or B-ALL patients and the survival curves were generated using Kaplan-Meier analysis.

**Supplemental Figure 16:**
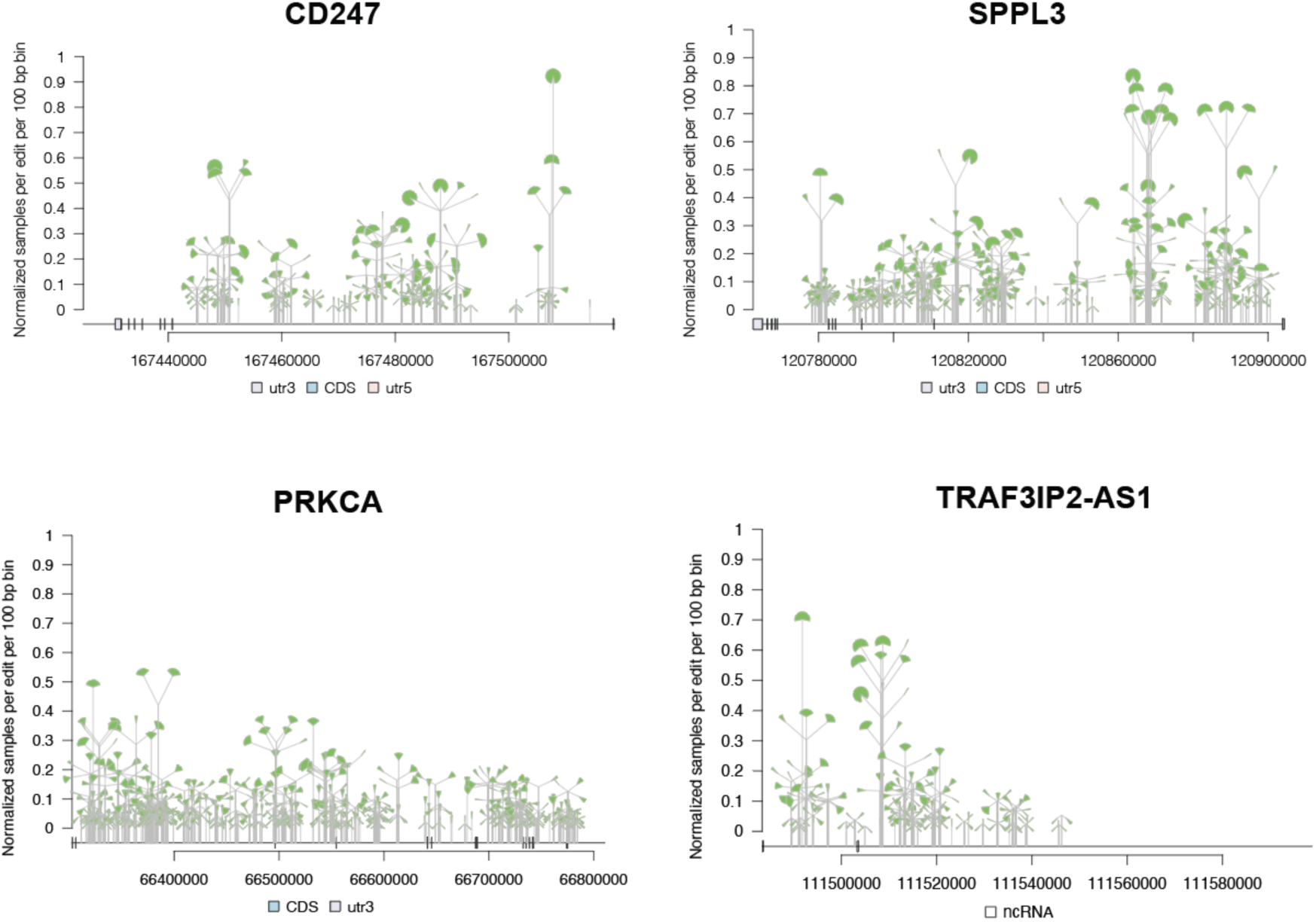
Dandelion plots showing the distribution of RNA editing sites within *CD247, SPPL3, PRKCA,* and *TRAF3IP2-AS1*. Each stem indicates the genomic position of an editing site, with the stem height representing the normalized editing signal in a smoothed 100 bp window, to help show where editing events cluster along the gene. At the top of each stem, a pie chart shows the fraction of 1,025 T-ALL or 37 B-ALL individuals that have an edit at the site. Gene models are shown below each plot.

